# Transcription apparatus of the yeast killer DNA plasmids: Architecture, function, and evolutionary origin

**DOI:** 10.1101/320143

**Authors:** Michal Sýkora, Martin Pospíšek, Josef Novák, Silvia Mrvová, Libor Krásný, Václav Vopálenský

## Abstract

Transcription of extrachromosomal elements such as organelles, viruses, and plasmids is dependent on cellular RNA polymerase (RNAP) or intrinsic RNAP encoded by these elements. The yeast *Kluyveromyces lactis* contains killer DNA plasmids that bear putative non-canonical RNAP genes. Here, we describe the architecture and evolutionary origin of this transcription machinery. We show that the two RNAP subunits interact *in vivo*, and this complex interacts with another two plasmid-encoded proteins - the mRNA capping enzyme, and a putative helicase which interacts with plasmid-specific DNA. Further, we identify a promoter element that causes 5’ polyadenylation of plasmid-specific transcripts *via* RNAP slippage during transcription initiation, and structural elements that precede the termination sites. As a result, we present a first model of the yeast killer plasmid transcription initiation and intrinsic termination. Finally, we demonstrate that plasmid RNAP and its promoters display high similarity to poxviral RNAP and promoters of early poxviral genes, respectively.

## INTRODUCTION

Linear double-stranded DNA plasmids were found in the cytoplasm of several yeast species. Structural organization of these plasmids is quite uniform and they are often present as two or three differently sized DNA elements in yeast host cells (Jeske et al. 2007). Characteristic features of these plasmids are terminal proteins covalently linked to the 5’ ends of their DNA, terminal inverted repeats, and their cytoplasmic localization (Gunge et al. 1982; Kikuchi et al. 1984; Stam et al. 1986). Yeast linear plasmids of *Kluyveromyces lactis*, termed pGKL1 (or K1) and pGKL2 (or K2), have become a model system to study such DNA elements. These plasmids have compact genomes with occasional overlaps of open reading frames (ORFs) and a high AT content of ~74% (Sor and Fukuhara 1985; Tommasino et al. 1988). The presence of both pGKL plasmids in several *K. lactis* strains is associated with the extensively studied yeast killer phenotype (Gunge et al. 1981).

Functions of protein products for most ORFs encoded by the pGKL plasmids were predicted using bioinformatics approaches and some of these proteins were characterized by biochemical and genetic analyses (Jeske et al. 2007). Both pGKL1 and pGKL2 encode their own DNA polymerase and a terminal protein, and it is assumed that the mechanism of their replication is similar to the replication of viruses of the *Adenoviridae* family or *Bacillus subtilis* bacteriophage φ29 (Stark et al. 1990). Consequently, marked sequence similarities between viral enzymes and putative products of several linear plasmid ORFs with expected function in replication and transcription resulted in yeast linear DNA plasmids being called virus-like elements (Satwika et al. 2012). Hence, it is believed that these linear plasmids may have originated from endosymbiotic bacteria or a virus (Kempken et al. 1992). Nevertheless, the exact evolutionary origin of the yeast linear plasmids remains unclear.

Transcription of plasmid-specific genes has been shown to be independent of mitochondrial (Gunge and Yamane 1984) and nuclear RNA polymerases (Romanos and Boyd 1988; Kämper et al. 1989; Stark et al. 1990; Kämper et al. 1991), and probably utilizes a plasmid-specific RNA polymerase (RNAP). Experiments with bacterial reporter and yeast nuclear genes fused with pGKL plasmid sequences identified an upstream conserved sequence (5’-ATNTGA-3’) preceding each of the open reading frames. This upstream conserved sequence (UCS), usually located at a distance of 20–40 nucleotides prior to the start codon, is essential for cytoplasmatic transcription of the downstream located gene (Schründer and Meinhardt 1995; Schickel et al. 1996; Schründer et al. 1996). Sequences located farther upstream of the UCS element have been shown to have no effect on transcription (Schickel et al. 1996). The UCS element is highly conserved among all yeast linear plasmids and the UCS sequence derived from the *Pichia etchellsii* pPE1B plasmid acts as a functional promoter when transplanted into the pGKL1 plasmid (Klassen et al. 2001). Thus, the UCS element is a universal cis-acting component of the plasmid-specific transcription system and it is essential for transcription initiation. After elongation, transcription then terminates after each gene because only monocistronic transcripts were revealed with Northern blot analyses of transcripts derived from ten pGKL-encoded ORFs (Romanos and Boyd 1988; Tommasino et al. 1988; Schaffrath et al. 1995b; Schaffrath et al. 1997; Jeske et al. 2006). This suggests the existence of a defined, yet unknown mechanism of transcription termination.

Unique RNAP subunits, and possibly also a putative helicase and the mRNA capping enzyme are the key elements of the plasmid cytoplasmic transcription machinery. Protein products of *ORF6* (K2ORF6p; large subunit) and *ORF7* (K2ORF7p; small subunit) of the plasmid pGKL2 should form a non-canonical RNAP. K2ORF6p was found to have a sequence similarity to three conserved regions of the two largest subunits (β and β’ in bacteria) of canonical multisubunit RNAPs (Wilson and Meacock 1988). Sequence similarity of K2ORF6p to the β and β’ subunit has recently been extended to 12 conserved regions shared by all bacterial, archaeal and eukaryotic RNAPs (Ruprich-Robert and Thuriaux 2010), and the predicted structure of this enzyme thus resembles a fusion of the β subunit with a portion of the β’ subunit. K2ORF7p was found to have sequence similarities to two conserved regions of the β’ subunit, which are usually located at the C-terminus of β’ (Schaffrath et al. 1995a).

The *ORF4* sequence of the plasmid pGKL2 shows striking sequence similarity to viral helicases from the superfamily II of DEAD/H family helicases involved in transcription. The *K2ORF4* protein product (K2ORF4p) displays similarity with two Vaccinia virus helicases – (i) NPH-I, which is encoded by the *D11L* gene, and (ii) the small subunit of the heterodimeric Vaccinia virus early transcription factor (VETF) encoded by the *D6R* gene (Wilson and Meacock 1988). NPH-I is known to provide the energy for elongation of transcription and for the release of RNA during transcription termination (Deng and Shuman 1998). VETF functions as a transcription initiation factor that binds and bends the promoter region of early genes (Broyles et al. 1991).

The protein product of *ORF3* (K2ORF3p) encoded by the plasmid pGKL2 shows sequence similarity to the Vaccinia virus mRNA capping enzyme encoded by the *D1R* gene that consists of three domains responsible for the three enzymatic activities necessary to form the 5’ mRNA cap structure (Larsen et al. 1998). The methyltransferase activity of the D1 protein of the poxvirus Vaccinia is allosterically stimulated by heterodimerization with a smaller protein encoded by the *D12L* gene (Mao and Shuman 1994; Schwer et al. 2006). The complex of D1 and D12 proteins is sometimes also referred to as the vaccinia termination factor (VTF) because, together with NPH-I, it also acts as a transcription termination factor of early genes (Luo et al. 1995). Triphosphatase and guanylyltransferase activities of K2ORF3p have been already confirmed experimentally *in vitro* (Tiggemann et al. 2001).

As reported previously, the *K2ORF3, K2ORF4, K2ORF6* and *K2ORF7* genes are indispensable for the maintenance of the pGKL plasmids in the cell (Schaffrath et al. 1995a, 1995b; Schaffrath et al. 1997; Tiggemann et al. 2001). However, understanding of interactions of their protein products with each other and with plasmid DNA in the cell is lacking, as well as understanding of linear plasmid DNA sequence elements required for transcription initiation and termination.

Here, we present a systematic *in vivo* study focusing on the architecture of the transcription complex of the yeast linear plasmids. Moreover, we identify a new promoter DNA element which is associated with 5’ mRNA polyadenylation of most pGKL-encoded genes and we uncover a link between RNA stem loop structures and 3’ end formation of plasmid-specific mRNAs *in vivo*. Further, we present an extensive phylogenetic analysis of amino acid sequences of linear plasmid RNAP subunits. Finally, we provide a detailed sequence analysis of pGKL promoters. Collectively, these analyses strongly suggest that the linear plasmid transcription machinery is closely related to the transcription machinery of poxviruses.

## RESULTS

### Plasmid RNAP subunits, mRNA capping enzyme, and helicase associate *in vivo*

To start characterizing the transcription machinery of the yeast linear plasmids we first tested whether the plasmid RNAP subunits (K2ORF6p, K2ORF7p), the mRNA capping enzyme (K2ORF3p), and the putative helicase (K2ORF4p) form a complex *in vivo*.

Initially, we tested interactions between K2ORF3p, K2ORF6p and K2ORF7p using a yeast two-hybrid system and its fluorescence variant called bimolecular fluorescence complementation but we failed to detect any interaction with either approach (data not shown). This was most likely caused by the high AT content of linear plasmid genes that was shown recently to impair their nuclear expression due to RNA fragmentation mediated by the polyadenylation machinery (Kast et al. 2015). Therefore, we decided to prepare modified pGKL plasmids expressing the putative transcription machinery components containing various tags.

Next, we prepared a strain encoding yeast enhanced green fluorescent protein 3 (yEGFP3) (Cormack et al. 1997) that was fused to the N-terminus of the large RNAP subunit K2ORF6p (yEGFP3-K2ORF6p; strain IFO1267_pRKL2–4). We immunoprecipitated (IP) yEGFP3-K2ORF6p from this strain using GFP-Trap_A agarose beads that contain a monoclonal antibody against common GFP variants. As a control, we used the wt strain IFO1267 that had no modifications. Extensive washing was used to remove weakly bound proteins. The bound proteins were eluted, resolved on SDS-PAGE, and stained with Coomassie Brilliant Blue G-250 (Figure 1). Gel lanes or bands of interest were excised from the gel, and analysed by mass spectrometry (MS). As shown in Table 1, peptides corresponding to yEGFP3-K2ORF6p, K2ORF7p (RNAP small subunit) and K2ORF3p (mRNA capping enzyme) were detected whereas no such peptides were identified in the parallel-treated IFO1267 control sample.

**Table 1.**
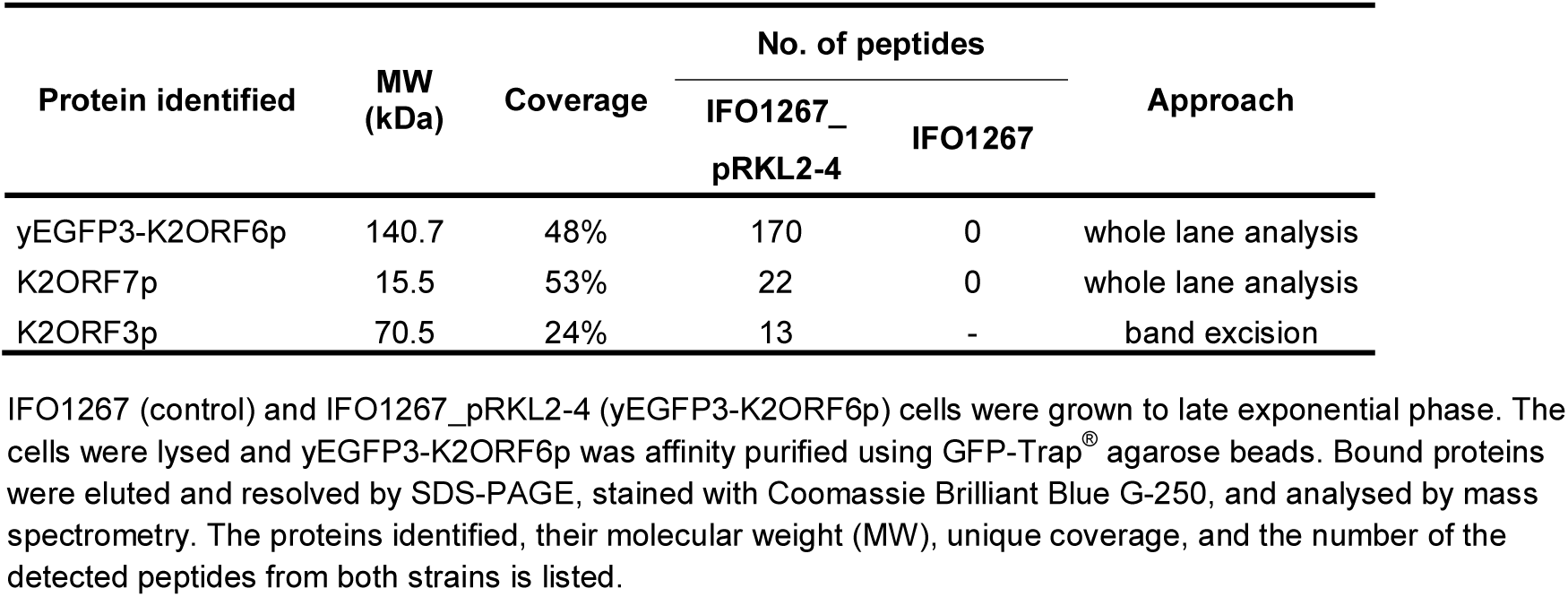
K2ORF6p-associated proteins.

| Protein identified | MW<br>(kDa) | Coverage | No. of peptides |  | Approach |
| --- | --- | --- | --- | --- | --- |
|  |  |  | IFO1267_<br>pRKL2-4 | IFO1267 |  |
| yEGFP3-K2ORF6p | 140.7 | 48% | 170 | 0 | whole lane analysis |
| K2ORF7p | 15.5 | 53% | 22 | 0 | whole lane analysis |
| K2ORF3p | 70.5 | 24% | 13 | - | band excision |

IFO1267 (control) and IFO1267\_pRKL2-4 (yEGFP3-K2ORF6p) cells were grown to late exponential phase. The cells were lysed and yEGFP3-K2ORF6p was affinity purified using GFP-Trap<sup>®</sup> agarose beads. Bound proteins were eluted and resolved by SDS-PAGE, stained with Coomassie Brilliant Blue G-250, and analysed by mass spectrometry. The proteins identified, their molecular weight (MW), unique coverage, and the number of the detected peptides from both strains is listed.

**Figure 1.**
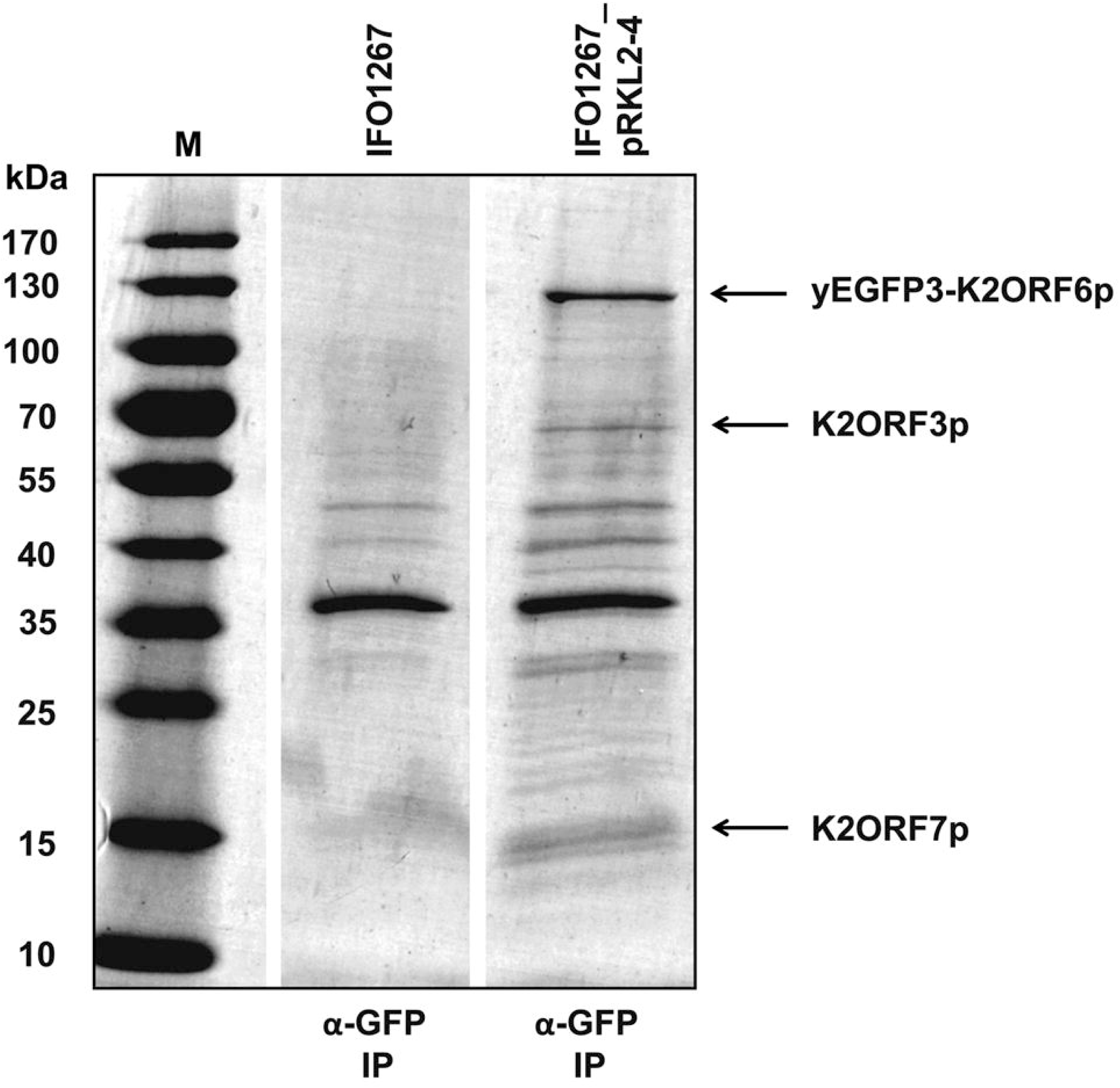
Identification of proteins associated with the large subunit of the linear plasmid RNAP (K2ORF6p). The gel shows Coomassie stained proteins affinity purified with GFP-Trap_A from strains IFO1267 (control) and IFO1267_pRKL2–4 (containing yEGFP3-K2ORF6p). Proteins identified by mass spectrometry are indicated with arrows on the right side, and also listed in Table 1. M, protein molecular mass marker (PageRuler™ Prestained Protein Ladder, Fermentas); the respective molecular mass values are indicated on the left side.

Then, to verify the interaction between the large RNAP subunit and the mRNA capping enzyme we decided to perform immunoprecipitation using tagged K2ORF3p as the bait. Hence, we prepared a strain encoding yEGFP3 fused to the C-terminus of K2ORF3p (IFO1267_pRKL2–11 strain). Interestingly, selective cultivation of clones after transformation led to a loss of the pGKL1 plasmid (Supplementary Figure S1). Immunoprecipitation using GFP-Trap_A agarose beads and subsequent MS analysis revealed peptides corresponding to K2ORF6p, K2ORF7p, K2ORF4p, and K2ORF3p-yEGFP3 (Supplementary Figure S2).

To further validate the results, we prepared several strains containing combinations of these proteins with various tags: (i) strain IFO1267_pRKL2–5 where yEGFP3-K2ORF6p was co-expressed together with K2ORF7p-Flag, and control strain IFO1267_pRKL2–15 that expressed only one tagged protein - K2ORF7p-Flag; (ii) strain IFO1267_pRKL2–6 coexpressing yEGFP3-K2ORF6p and K2ORF3p-HA, and control strain IFO1267_pRKL2–14 expressing only K2ORF3p-HA. The results of IP experiments followed with Western blotting clearly showed specific interactions between these proteins (Figures 2A and 2C).

**Figure 2.**
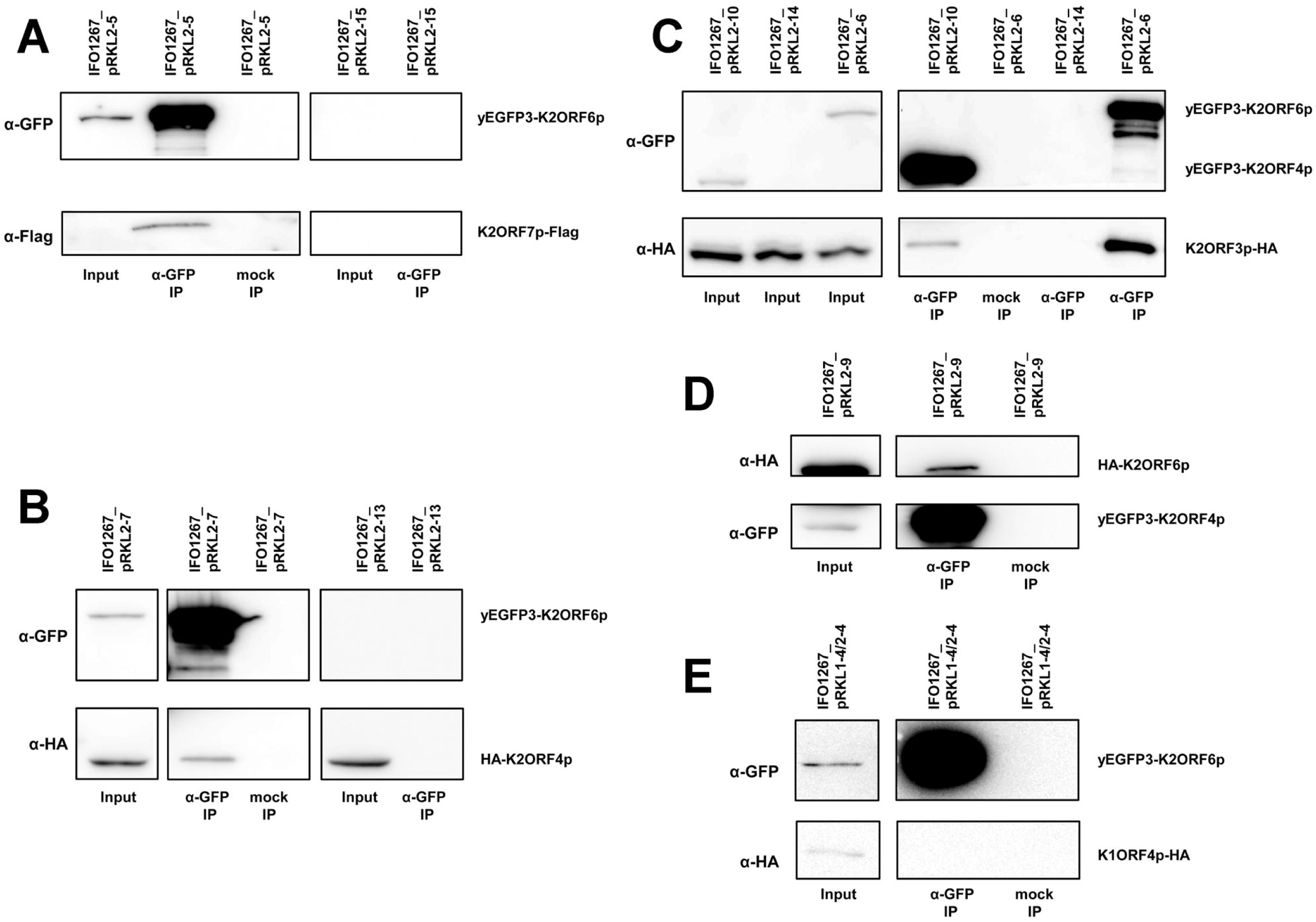
Interactions between RNAP subunits, mRNA capping enzyme, and putative helicase of the yeast linear plasmids. **(A)** Western blot of immunoprecipitated (α-GFP IP) and mock immunoprecipitated (mock IP) proteins from IFO1267_pRKL2–5 (yEGFP3-K2ORF6p, K2ORF7p-Flag) or control IFO1267_pRKL2–15 cells (K2ORF7p-Flag), respectively. The strains used are indicated above the lanes. The antibodies used for Western blot are indicated on the left hand side of the strips. The proteins detected are indicated on the right hand side of the strips. Positions of the identified proteins corresponded with theoretical molecular weight of the full length recombinant proteins, as determined by positions of the protein mass markers (not shown). Input represented approximately 1/100 of the sample and IP represented approximately 1/2 of the sample in this and the other immunoprecipitation experiments. Mock immunoprecipitations in all experiments were done using empty agarose beads. The same experimental scheme is used throughout this figure. (**B**) Western blot analysis of immunoprecipitations from lysates from IFO1267_pRKL2–7 (yEGFP3-K2ORF6p, HA-K2ORF4p) and control IFO1267_pRKL2–13 (HA-K2ORF4p) cells (indicated above the lanes). The (α-GFP) and anti-HA (α-HA) antibodies used are indicated on the left hand side; the detected proteins on the right hand side. (**C**) Western blot analysis of immunoprecipitations from lysates from strains IFO1267_pRKL2–6 (yEGFP3-K2ORF6p, K2ORF3p-HA), IFO1267_pRKL2–10 (yEGFP3-K2ORF4p, K2ORF3p-HA), and IFO1267_pRKL2–14 (control). (**D**) Western blot analysis of immunoprecipitations from lysates from IFO1267_pRKL2–9 cells (yEGFP3-K2ORF4p, HA-K2ORF6p). (**E**) Western blot analysis of immunoprecipitations from lysates from IFO1267_pRKL1–4/2–4 cells (yEGFP3-K2ORF6p, K1ORF4p-HA).

Thus far, we had detected K2ORF3p, K2ORF6p, and K2ORF7p to be associated *in vivo*. For the fourth protein, the putative helicase K2ORF4p, the MS results suggesting it as part of the complex were not convincing due to low protein coverage (Supplementary Figure S2).To address whether it does, although perhaps weakly, interact with these proteins, we prepared a strain expressing HA-K2ORF4p together with yEGFP3-K2ORF6p (IFO1267_pRKL2–7), and a control strain expressing HA-K2ORF4p only (IFO1267_pRKL2–13). After IP and Western blotting we found HA-K2ORF4p to associate with yEGFP3-K2ORF6p (Figure 2B). The results clearly showed that the putative helicase was specifically associated with the large RNAP subunit (or, was present in a complex containing this subunit) because yEGFP3-K2ORF6p and HA-K2ORF4p were not bound to the empty agarose beads, and HA-K2ORF4p alone did not bind to the GFP-Trap antibody (Figure 2B).

Because the association of the putative helicase with the large RNAP subunit seemed rather weak, we decided to perform reciprocal immunoprecipitation. Further, we also tested whether the mRNA capping enzyme associated with the putative helicase. Strains expressing (i) yEGFP3-K2ORF4p together with HA-K2ORF6p (IFO1267_pRKL2–9), and (ii) yEGFP3-K2ORF4p together with K2ORF3p-HA (IFO1267_pRKL2–10) were prepared. With the first combination we confirmed that HA-K2ORF6p associated with yEGFP3-K2ORF4p (Figure 2D). With the second combination we found that K2ORF3p-HA associated with yEGFP3-K2ORF4p (Figure 2C). The detected interactions were specific because yEGFP3-K2ORF4p and HA-K2ORF6p were not bound to the empty agarose beads and K2ORF3p-HA alone did not bind to the GFP-Trap antibody (Figures 2C and 2D).

As an additional control to demonstrate that the observed interactions were specific, we used another pGKL-encoded protein with a function unrelated to transcription. We selected K1ORF4p, a subunit of the toxin responsible for plasmid-associated killer yeast phenotype (Tokunaga et al. 1989). A strain co-expressing yEGFP3-K2ORF6p together with K1ORF4p-HA (IFO1267_pRKL1–4/2–4) was prepared. We found that K1ORF4p-HA was not associated with yEGFP3-K2ORF6p (Figure 2E).

Finally, we wanted to know whether the association of the putative helicase and mRNA capping enzyme with the large RNAP subunit was dependent on nucleic acids. We prepared lysates from IFO1267_pRKL2–6 (yEGFP3-K2ORF6p, K2ORF3p-HA) and IFO1267_pRKL2–7 (yEGFP3-K2ORF6p, HA-K2ORF4p) strains. The lysates were incubated with GFP-Trap_A beads and, after washing, the beads were split into two parts which were treated or mock-treated with Benzonase Nuclease to digest DNA and RNA. Then, the beads were again extensively washed and the bound proteins were eluted. Subsequent Western blot analysis revealed both K2ORF3p-HA and HA-K2ORF4p to associate with yEGFP3-K2ORF6p regardless of the presence or absence of nucleic acids which was confirmed by PCR (Supplementary Figure S3).

Taken together, the immunoprecipitation, mass spectrometry, and Western blot results demonstrated the existence of the hypothesized plasmid-specific transcription complex because K2ORF3p, K2ORF4p, K2ORF6p, and K2ORF7p were specifically associated *in vivo*. This association was independent of nucleic acids. Finally, K2ORF3p, K2ORF6p, and K2ORF7p appeared to form a core transcription complex with relatively strong mutual interactions to which K2ORF4p bound relatively weakly.

### Putative helicase is associated with plasmid-specific DNA *in vivo*

The weak binding of K2ORF4p to the other three proteins suggested that the putative helicase may also interact with plasmid DNA. Hence, we performed *in vivo* chromatin immunoprecipitation. We used the IFO1267_pRKL2–13 strain expressing HA-K2ORF4p and the IFO1267 control strain. First, we verified that the mouse monoclonal anti-HA HA-7 agarose efficiently immunoprecipitated HA-K2ORF4p (Figure 3A). Then, we performed chromatin immunoprecipitation of HA-K2ORF4p from formaldehyde cross-linked cells using mouse monoclonal anti-HA HA-7 agarose. The immunoprecipitated and input DNAs were used as templates for subsequent PCR analysis using primers designed to detect chromosomal or linear plasmid DNA. We used primers specific for *K. lactis* actin *(ACT)* and high-affinity glucose transporter *(HGT1)* genes as markers of chromosomal DNA, and toxin immunity *(K1ORF3)* and mRNA capping enzyme *(K2ORF3)* genes as markers of pGKL plasmids. We found, that HA-K2ORF4p was specifically associated with pGKL plasmids and not with chromosomal DNA (Figure 3B). These results were also confirmed by semiquantitative real-time PCR (data not shown).

**Figure 3.**
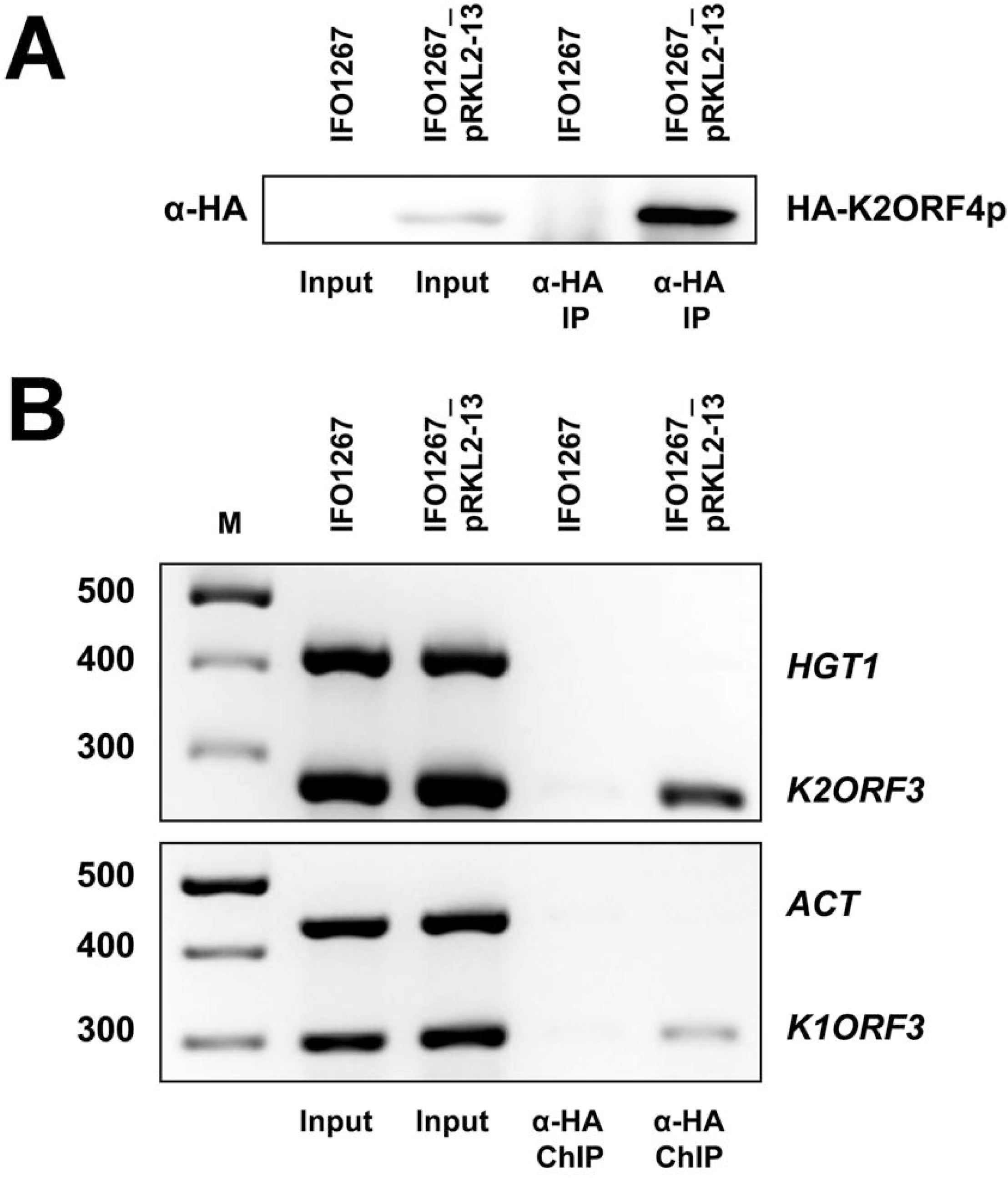
Physical association of the putative helicase (K2ORF4p) of the yeast linear plasmids with plasmid-specific DNA. (**A**) Western blot of HA-K2ORF4p that was affinity purified from lysates of IFO1267_pRKL2–13 (HA-K2ORF4p) and IFO1267 (control) cells. The strains used are indicated above the lanes. The antibody used is indicated on the left hand side of the strip. The protein detected is indicated on the right hand side of the strip. (**B**) PCR analysis of the presence of chromosomal *(ACT, HGT1)* or linear plasmid *(K1ORF3, K2ORF3)* DNA in chromatin immunoprecipitated using anti-HA HA-7 agarose from IFO1267_pRKL2–13 (HA-K2ORF4p) and IFO1267 (control) cells. Samples of individually performed gene-specific PCRs were analysed in 2.5 % agarose gel stained with ethidium bromide. The identity of the bands (genes) is indicated on the right. M, DNA molecular mass marker (GeneRuler™ 100 bp Plus DNA Ladder, Fermentas). The respective values are indicated on the left.

We concluded that the previously uncharacterized helicase was associated with plasmid-specific DNA *in vivo*. Due to relatively weak interaction of the putative helicase with the core of plasmid-specific transcription complex *in vivo*, we assume that it possibly acts as a dissociable transcription factor.

### Slippage of RNAP at the initiation site results in 5’ polyadenylation of plasmid mRNAs

Next, we wished to characterize transcription initiation of the linear plasmid genes. Our previous 5’ RACE-PCR experiments had revealed 5’ cap structures on the plasmid-specific mRNAs, likely synthetized by plasmid-encoded K2ORF3p mRNA capping enzyme, and also the presence of non-templated 5 poly(A) leaders of heterogeneous lengths in mRNAs of 12 pGKL genes except for *K2ORF2, K2ORF3* and *K2ORF8* (manuscript submitted). Interestingly, heterogeneous 5 poly(A) leaders are a known feature of poxviral intermediate and late transcripts (Bertholet et al. 1987; Schwer et al. 1987). It was shown that the 5’ poly(A) leader was produced by slippage of Vaccinia virus RNAP on three consecutive thymidine residues in the template strand at the initiator region (INR) where transcription starts both *in vivo* and *in vitro* (Schwer and Stunnenberg 1988; Davison and Moss 1989).

Figures 4A and 4B show the sequence logo of the INR consensus motif (TAAAT) of Vaccinia virus intermediate and late genes, respectively (Yang et al. 2012). Interestingly, we were able to locate a similar INR-like consensus sequence (TAAAN) in promoters of all 12 pGKL genes whose transcripts were 5’ polyadenylated (Figure 4C). The first adenosine residue of the motif was considered to be the initiating nucleotide for the TSS annotation.

**Figure 4.**
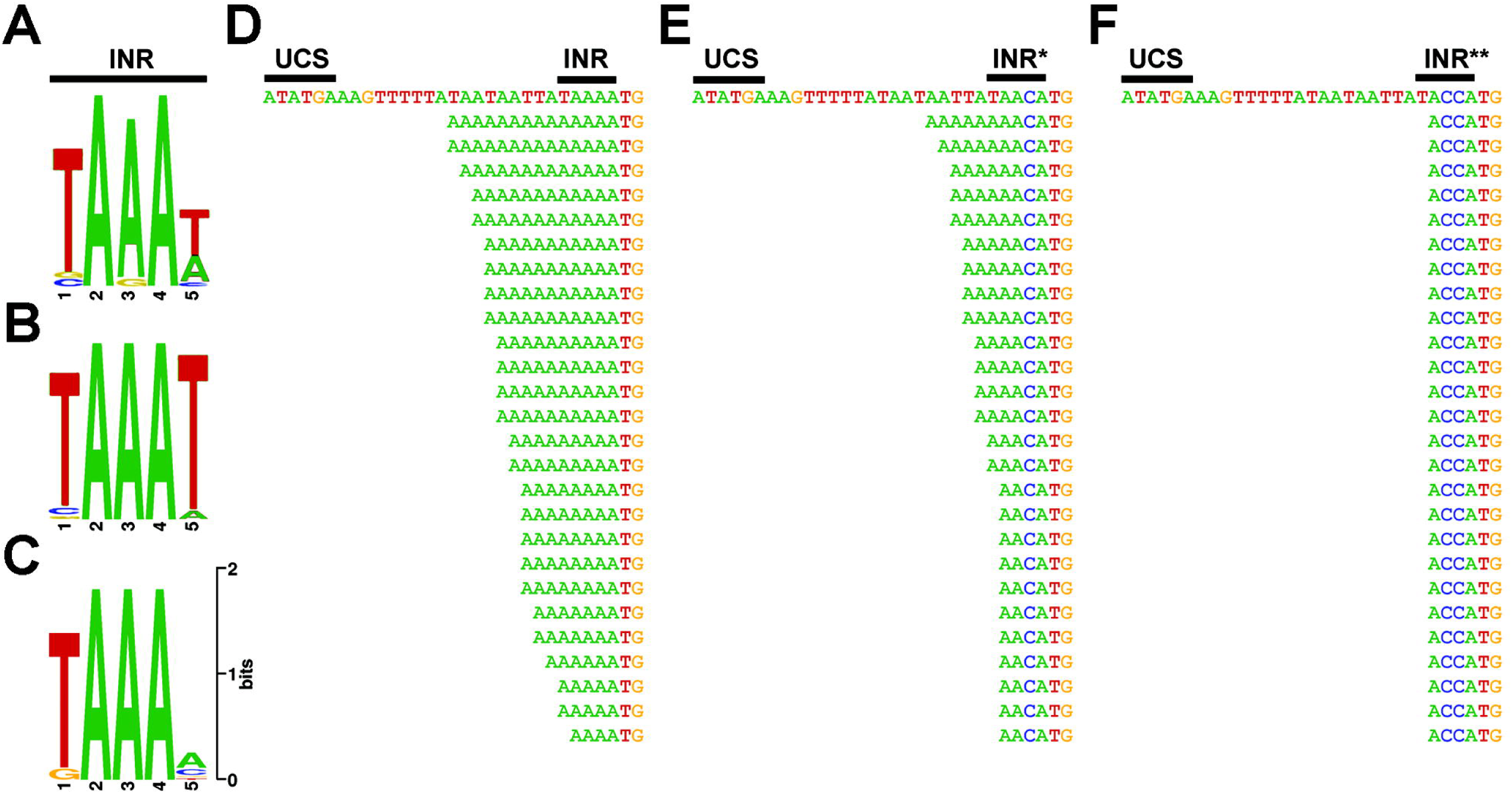
Promoters of yeast linear plasmids contain initiator region (INR) responsible for non-templated 5’ polyadenylation of mRNAs. (**A**) Sequence logo of the initiator region in promoter sequences of Vaccinia virus intermediate genes (Yang et al. 2012). (**B**) Sequence logo of the initiator region in promoter sequences of Vaccinia virus late genes (Yang et al. 2012). (**C**) Sequence logo of the putative INR identified in promoters of 12 ORFs encoded by pGKL plasmids. (**D**) 5’ RACE-PCR analysis of the *G418^R^* gene from the IFO1267_pRKL1–1 strain. In this and the following panels, the upper sequence corresponds to the template (plasmid) DNA and the UCS is indicated; sequences situated below represent individual sequenced cDNA clones (the 5 untranslated region is shown in full till the translation start codon, ATG). Guanosine residues corresponding to the original 5’ mRNA caps which were present in some of the cDNA clones are omitted in this representation for clarity. (**E**) 5’ RACE-PCR analysis of the *G418^R^* gene from the IFO1267_pRKL1–2 strain bearing a promoter mutation in the putative INR reducing the number of consecutive adenosine residues in the template. (**F**) 5’ RACE-PCR analysis of the *G418^R^* gene from the IFO1267_pRKL1–3 strain bearing promoter mutations in the putative INR abolishing consecutive adenosine residues in the template.

Subsequently, we tested whether the putative INR was responsible for the 5 end polyadenylation of the pGKL-derived transcripts. We prepared three *K. lactis* strains with modified pGKL1 plasmids encoding the G418 resistance marker under the control of the K1UCR2 promoter. We prepared three variants of the K1UCR2 promoter that differed in the INR sequence: (i) TAAAA (wt; strain IFO1267_pRKL1–1); (ii) TAACA (strain IFO1267_pRKL1–2); and (iii) TACCA (strain IFO1267_pRKL1–3). Then, we purified total RNA from the three strains, prepared cDNA, and performed 5 RACE-PCR to determine the 5’ end sequences. The results showed that the 5’ poly(A) leader was present when the K1UCR2 sequence contained the putative wt INR (TAAAA INR) (Figure 4D) When TAACA was used, the length of the 5’ end poly(A) was significantly reduced (Figure 4E). When TACCA was used, the poly(A) leader disappeared altogether (Figure 4F).

Therefore, we concluded that slippage of plasmid RNAP at the initiation site was the mechanism responsible for 5’ polyadenylation of the transcripts. Moreover, the identified INR sequence constituted an independent DNA element, not influenced by the sequence of the gene because the pattern of the sequenced 5’ RACE-PCR clones for *K1ORF2* transcripts was the same as for *G418^R^* transcripts produced from the K1UCR2 with the wt INR (manuscript submitted).

### RNA stem loop structures influence 3’ end formation of plasmid-specific mRNAs *in vivo*

Our previous 3’ RACE-PCR experiments had revealed the absence of 3’ poly(A) tails in mRNAs of all 15 pGKL ORFs (manuscript submitted). To shed light on the mode of transcription termination of the linear plasmids, we tried to identify sequence/secondary structure elements/signals near the 3’ termini. First, we searched for sequence motifs within the last 150 nt of each transcript that would be shared among the 15 pGKL ORFs but we detected none. Second, we searched for secondary structure motifs using the RNAstructure Server (Reuter and Mathews 2010). We identified putative RNA stem loop structures close to the experimentally determined 3’ ends of cDNA (Supplementary Figure S4). The putative RNA stem loops were typically in the vicinity of the respective ORF’s stop codon with the median distance of 26 nt, and Gibbs free energy of −7.5 kcal/mol.

Hence, we tested, whether the predicted RNA stem loop structures influenced the 3 mRNA end formation. Because pGKL plasmids contain almost no intergenic regions, the putative RNA stem loops are localized in the coding sequences of adjacent ORFs or within the terminal inverted repeats. This means that their sequences cannot be subjected to mutagenesis without the possibility of altering plasmid functions. Therefore, we prepared a *K. lactis* strain with a modified pGKL1 plasmid encoding the G418 resistance marker under control of K1UCR2, followed by the 3’ UTR of the *K2ORF5* gene (strain IFO1267_pRKL1–5; Figure 5A). The distal part of the *K2ORF5* 3’ UTR contained two putative partially overlapping RNA stem loops termed Stem loop 1 and 2 (Supplementary Figure S4I). 3 RACE-PCR experiments revealed a transcription termination pattern that could be attributed to the presence of both Stem loop 1 and 2 (Figures 5B and 5C). Next, using the same promoter-gene-3’ UTR arrangement, we prepared a strain with 4 nucleotide mutations destabilizing the base pairing in the middle of the putative Stem loop 2 (strain IFO1267_pRKL1–6). For this construct, we detected a transcription termination pattern that could be attributed to the presence of only Stem loop 1 (Figures 5B and 5D). Subsequently, we prepared a strain (IFO1267_pRKL1–7) where we changed the sequence but not the base-pairing of 4 nucleotides within the Stem loop 2. 3’ RACE-PCR experiments revealed a transcription termination pattern that could be attributed to the presence of the rescued Stem loop 2 (Figures 5B and 5E). Note that the rescue mutations distinctly altered the length and the Gibbs free energy (destabilizing Stem loop 1) of the overlapping Stem loop 1 and this was likely the reason why transcription termination from the Stem loop 1 was not detected in the IFO1267_pRKL1–7 strain (Figures 5B and 5E).

**Figure 5.**
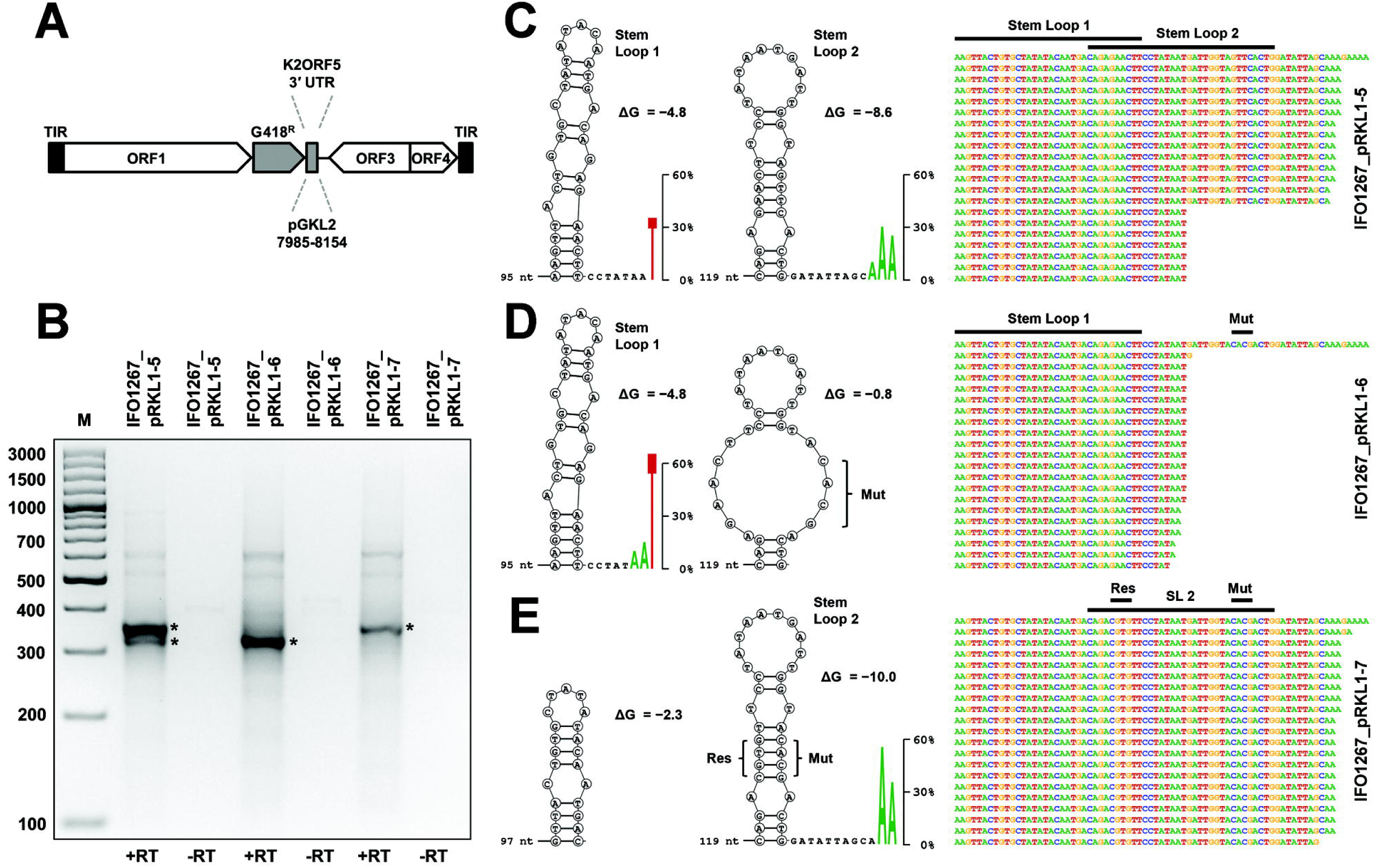
RNA stem loop structures influence the 3’ end formation of plasmid-specific mRNAs *in vivo*. (**A**) Schematic representation of recombinant pGKL1 plasmids where the *G418^R^* gene is followed by the coding sequence of wild-type (pRKL1–5) or modified (pRKL1–6, pRKL1–7) 3’ untranslated region of the *K2ORF5* gene. TIR - terminal inverted repeat. (B) 3’ RACE-PCR analysis of individual mRNAs corresponding to the *G418^R^* gene expressed from modified pGKL1 plasmids. Samples were analyzed in 3.0% agarose gel stained by ethidium bromide. The strains used to purify the RNA are indicated above the lanes. M, DNA molecular mass marker (GeneRuler™ 100 bp Plus DNA Ladder, Fermentas). The respective values are indicated on the left. Specific products that were cloned to the pCR^TM^4-TOPO^®^ vector and used for sequencing are labelled with asterisks. Reverse transcription was carried out in the presence (+RT) and absence (−RT) of reverse transcriptase. (**C**) 3’ RACE-PCR analysis of the *G418^R^* gene from IFO1267_pRKL1–5 strain. In this and the following panels, the upper sequences on the right correspond to the template (plasmid) DNA; sequences situated below represent 3’ end regions of individual sequenced cDNA clones. Positions of the putative RNA stem loops are indicated above the sequences. Predicted RNA stem loops are displayed as cDNA nucleotide letters in circles on the left and the values of Gibbs free energy (ΔG) in kcal/mol are displayed for each structure. Stem loop distances from the gene stop codon are shown as numbers of nucleotides. The last few 3’ end nucleotides of the experimentally determined 3’ ends of cDNA are shown as letters enlarged proportionally to their occurrence (in %) in the sequenced clones in the case when these nucleotides were detected in at least two independent clones. (**D**) 3’ RACE-PCR analysis of the *G418^R^* gene from the IFO1267_pRKL1–6 strain. Mut, the mutated stem loop. (**E**) 3’ RACE-PCR analysis of the *G418^R^* gene from IFO1267_pRKL1–7 strain. Mut, the mutated stem loop; Res, the rescued stem loop.

Finally, we determined whether promoter sequences or coding sequences of a gene could affect the 3’ mRNA end formation. We used *K2ORF5*, G418, and hygromycin B resistance genes under the control of K2UCR5, K1UCR2 and K1UCR3, respectively, located on the pGKL2 plasmid. Downstream of the coding sequence of each of these genes we positioned the 3 untranslated region (UTR) of the *K2ORF5* gene. We purified total RNA from the respective *K. lactis* strains, prepared cDNA, and performed 3’ RACE-PCR experiments. Although the coding sequences of the aforementioned genes differed both in length and AT content, the pattern of their 3’ termini was highly similar (Supplementary Figures S4 and S5).

We concluded that RNA stem loop structures were essential for the 3 end formation of plasmid-specific mRNAs *in vivo*, presumably acting as factor-independent intrinsic terminators. Moreover, this termination was independent of the promoter and the gene used both with respect to its sequence and length.

### Plasmid-specific RNA polymerase has unique architecture

To facilitate interpretation of the experimental data we created a 3D model of the pGKL-specific linear plasmid RNAP. This was feasible due to sequence similarity between parts of K2ORF6p, K2ORF7p and conserved regions of the canonical multisubunit RNAPs (Schaffrath et al. 1997; Ruprich-Robert and Thuriaux 2010). The structural models of the two subunits covered 92.7% of the K2ORF6p sequence and 62.1% of the K2ORF7p sequence, respectively.

Superimposition of the models over the *S. cerevisiae* RNAP II elongation complex (PDB ID: 2NVQ) showed that all known regions of K2ORF6p and K2ORF7p with sequence similarity to conserved regions of the canonical RNAPs were modelled accordingly. Interestingly, parts of K2ORF6p were modelled by βa1, βa6, βa13, and βa16 conserved regions, which were not detected previously to be present in K2ORF6p. We verified the proper model:template alignment of these regions by manually constructed sequence alignments with other canonical RNAPs and these alignments indeed showed sequence similarities between K2ORF6p and the aforementioned regions (Supplementary Figure S6).

The overall distribution of the conserved regions within K2ORF6p and K2ORF7p is depicted in Figure 6A. It should be noted that K2ORF6p displayed a unique fusion between and β’ subunit conserved regions, which is not known to be present in any other canonical or non-canonical RNAP. This fusion seemed to be essential for the maintenance of pGKL plasmids in yeast - a plasmid where we divided *K2ORF6* into two genes, based on their homology to β and β’, was unable to substitute wt pGKL2 in the cell (data not shown). On the other hand, a division of the β’ subunit (between β’a15 and β’a16) into two polypeptides, similarly as in K2ORF6p and K2ORF7p, is present in RNAPs of some *Archaeal* species (Kwapisz et al. 2008).

**Figure 6.**
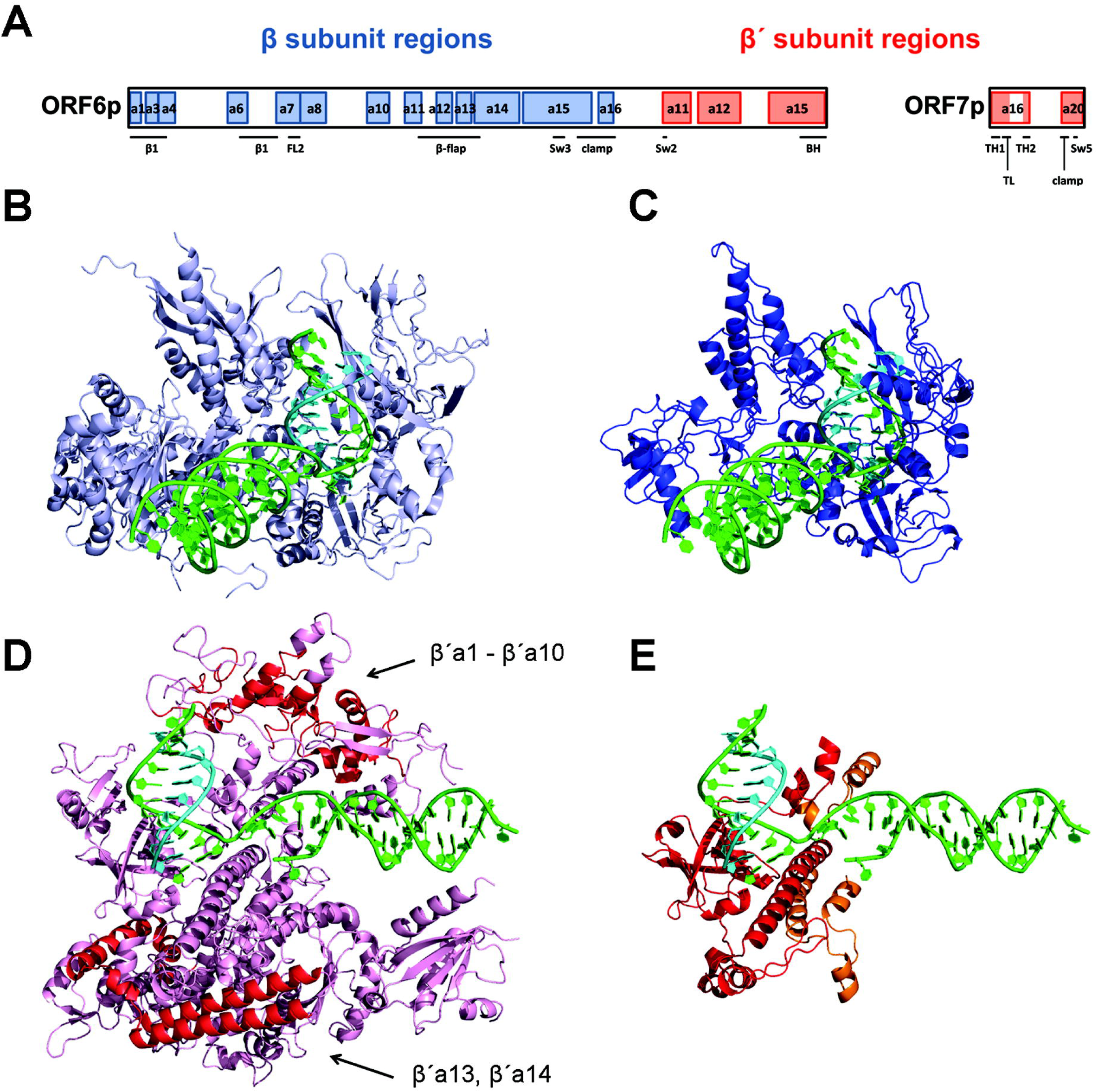
Predicted 3D structure of yeast linear plasmid non-canonical RNAP encoded by the *ORF6* and *ORF7* genes. (**A**) Schematic representation of the primary sequence of pGKL-encoded RNAP showing similarity to conserved regions of the catalytic subunits of canonical multisubunit RNAPs. β subunit conserved regions (blue) and β’ subunit conserved regions (red) present in K2ORF6p and K2ORF7p are drawn to scale. Conserved regions are named according to ref. (Lane and Darst 2010). Sequence alignments for newly detected similarity of *ORF6* protein products to βa1, βa6, βa13 and βa16 conserved regions are provided in Supplementary Figure S6. BH, Bridge helix; FL2, Fork loop 2; Sw2, Switch 2; Sw3, Switch 3; Sw5, Switch 5; TH1, Trigger helix 1; TH2, Trigger helix 2; TL, Trigger loop. (**B**) 3D crystal structure of *Saccharomyces cerevisiae* RNA polymerase II elongation complex showing the Rpb2 subunit (β subunit homolog, light blue), DNA (green) and RNA (cyan). This figure is based on 2NVQ. (**C**) 3D model of pGKL RNAP showing K2ORF6p residues 1–693 (β’ subunit homolog, blue), DNA (green) and RNA (cyan). Nucleic acids in this as well as the following structures are based on 2NVQ. (**D**) 3D crystal structure of *Saccharomyces cerevisiae* RNAP II elongation complex showing the Rpb1 subunit (β’ subunit homolog, pink), DNA (green) and RNA (cyan). Arrows indicate β’ regions shared by multisubunit RNAPs (red) that are clearly missing in the linear plasmid RNAP. This figure is based on 2NVQ. (**E**) 3D model of pGKL RNAP showing K2ORF6p residues 754–882 and 894–974 (β’ subunit homolog, red), K2ORF7p residues 1–52 and 103–132 (β’ subunit homolog, orange), DNA (green) and RNA (cyan). For details concerning structure modelling see Supplementary Materials and Methods.

Figures 6B and 6C show the *S. cerevisiae* RNAP II Rpb2 subunit (β subunit homolog) and the K2ORF6p model, respectively. It is clear that almost all conserved β subunit regions are present in K2ORF6p, and only the spacing between them is shorter. Figures 6D and 6E show the *S. cerevisiae* RNAP II Rpb1 subunit (β’ subunit homolog) and relevant homologous portions of K2ORF6p/K2ORF7p subunits, respectively. Remarkably, more than half of the β’ subunit conserved regions, such as most of the clamp domain (β’a1 - β’a10 regions) and secondary-channel rim helices (β’a13, β’a14 regions), are missing in plasmid RNAP Figure 6D.

We concluded that the linear plasmid RNAP displayed a unique and novel architecture.

### Plasmid-specific RNA polymerase has a viral origin

An extensive phylogenetic analysis of yeast linear plasmid RNAPs has not been performed yet. Earlier analyses focused only on conserved regions β’a11 and β’a12 of K2ORF6p. These studies suggested that plasmid RNAP was closer to multisubunit RNAPs rather than to single-subunit RNAPs encoded by mitochondrial linear plasmids of fungi and plants (Kempken et al. 1992; Rohe et al. 1992). Hence, it is believed that the *ORF6* and *ORF7* genes of linear plasmids were derived from eukaryotic RNAP genes of ancestral host cells (Jeske et al. 2007; Ruprich-Robert and Thuriaux 2010) or, alternatively, that the *ORF6* gene is an ancient representative of multisubunit RNAP diversification from times when and constituted a single protein (Iyer and Aravind 2012).

We performed a detailed phylogenetic analysis to delve deep into the evolutionary past of yeast linear plasmids. We used sequences of βa1, βa3-βa4, βa6-βa8, βa10-βa16, β’a11-β’a12, β’a15-β’a16 and β’a20 conserved regions of ORF6 and ORF7 proteins from all sequenced yeast linear plasmids. These conserved regions were divided into two alignments corresponding to β and β’ subunit regions. These alignments were then combined with the published alignments of β and β’ subunit conserved regions of canonical multisubunit RNAPs (Lane and Darst 2010), and they were used for construction of phylogenetic trees.

Figures 7A and 7B show the phylogenetic trees for and subunit conserved regions, respectively. The trees show that plasmid RNAPs branch together with viral RNAPs of the *Poxviridae* family, and plasmid RNAP β’ subunit regions are also branching close to viral RNAPs of the *Iridoviridae* family (Figure 7B). This is consistent with the prevalent view that original β’ subunit orthologs of all nucleo-cytoplasmic viruses are monophyletic, derived from eukaryotic RNAP I (Yutin and Koonin 2012).

**Figure 7.**
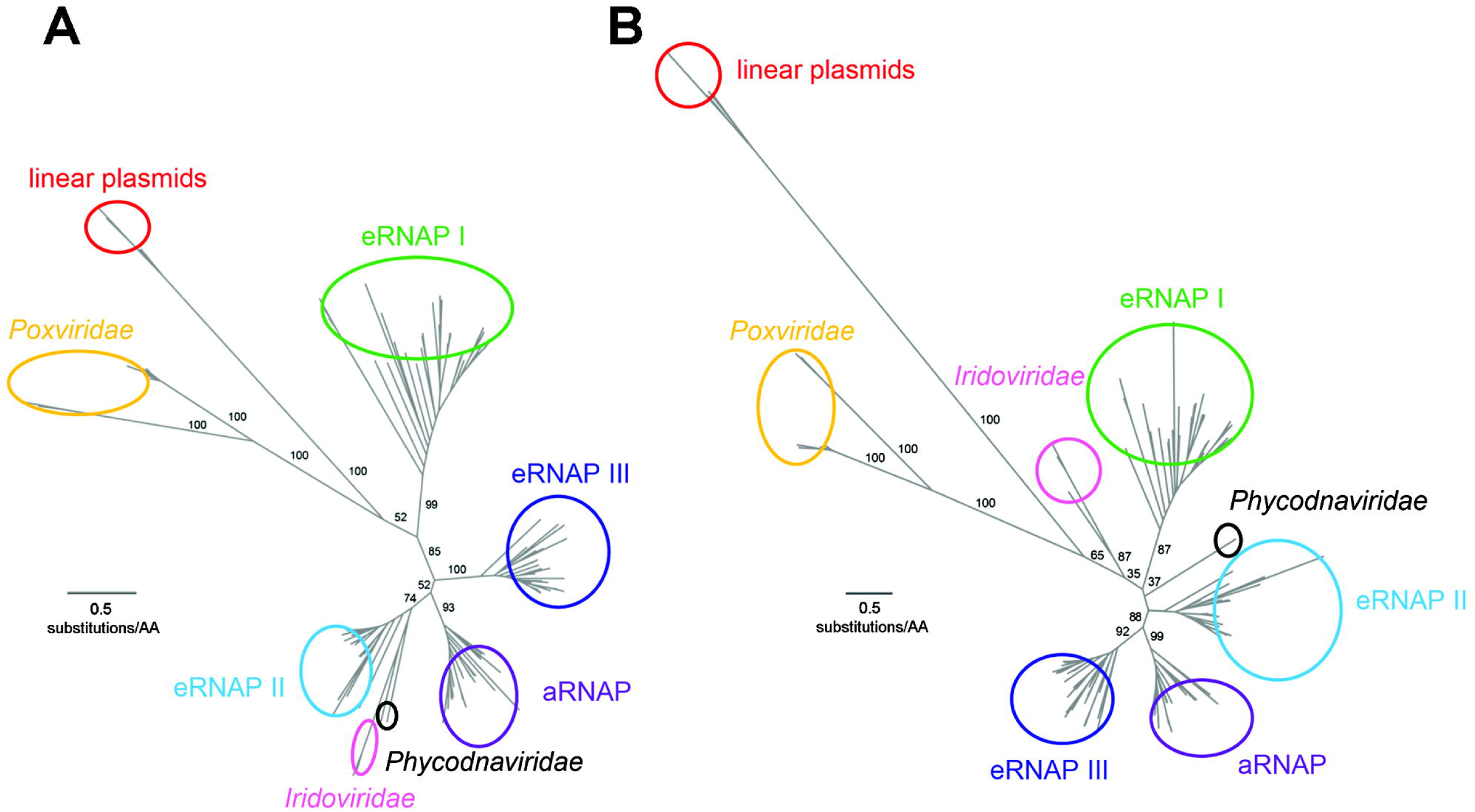
Phylogenetic analysis of yeast linear plasmid RNA polymerases with selected canonical RNAPs. Phylogenetic trees are displayed as unrooted phylograms where the length of the branches is proportional to the calculated evolutionary distance of individual sequences. Leaves defining the different classes of RNAPs are labeled. Used abbreviations for RNAP groups: aRNAP, archaeal RNAP; eRNAP, eukaryotic RNAP. Length scale of branches of an average value of 0.5 substitutions per amino acid residue is shown as a line near its corresponding tree. Selected branch support values calculated from 1000 bootstrap replicates are indicated in black. (**A**) Phylogram of β subunit homologs based on amino acid sequence alignment of β subunit conserved regions of selected canonical RNAPs and those β subunit conserved regions present in *ORF6* genes of the yeast linear plasmids. (**B**) Phylogram of β’ subunit homologs based on amino acid sequence alignment of subunit conserved regions of selected canonical RNAPs and those β’ subunit conserved regions present in *ORF6* and *ORF7* genes of the yeast linear plasmids. For details concerning phylogenetic analysis see Supplementary Materials and Methods.

Taken together, our phylogenetic analysis surprisingly points to a viral origin of plasmid RNAPs close to poxviruses, which is in contradiction to all previous hypotheses about the origin of these enzymes.

### Plasmid promoters have a viral origin

Although the UCS (5’-ATNTGA-3’) essential for linear plasmid transcription was identified a long time ago, no similarities with known promoters that would indicate its origin were reported. We extended the UCSs preceding all pGKL-encoded ORFs both upstream and downstream by ~10 bp, and we created a consensus motif. This consensus was then used to search for similar elements that were associated with transcription by multisubunit RNAPs. We particularly focused on promoters of viral RNAPs because our phylogenetic analysis of plasmid RNAPs had suggested a viral origin.

Notably, we detected great sequence similarity between the extended UCS (Figure 8C) and the upstream control element (UCE), which is a promoter element of poxviral early genes (Figure 8A). The UCE motif is a 15-nt long AT-rich element with any nucleotide at the 5^th^ position followed by TGA (Yang et al. 2011). This perfectly matched the extended UCS motif. The median distances from the 3’ ends of the UCEs to the annotated TSSs of 84 Vaccinia virus ORFs displayed a median distance of 12 nt (Figure 8B) (Yang et al. 2011). We annotated the TSSs of all pGKL plasmid genes based on our previous 5’ RACE-PCR experiments (Supplementary Materials and Methods, and Supplementary Table S4). The distances from the extended UCSs to the annotated TSSs of 15 pGKL-encoded ORFs Figure 8D had a median distance of 11 nt. Moreover, we found an adenosine residue to be the TSS nucleotide in all pGKL-encoded ORFs (Supplementary Table S4), similar to the TSSs of the poxviral early genes where purines are the dominant TSS bases (Yang et al. 2011).

**Figure 8.**
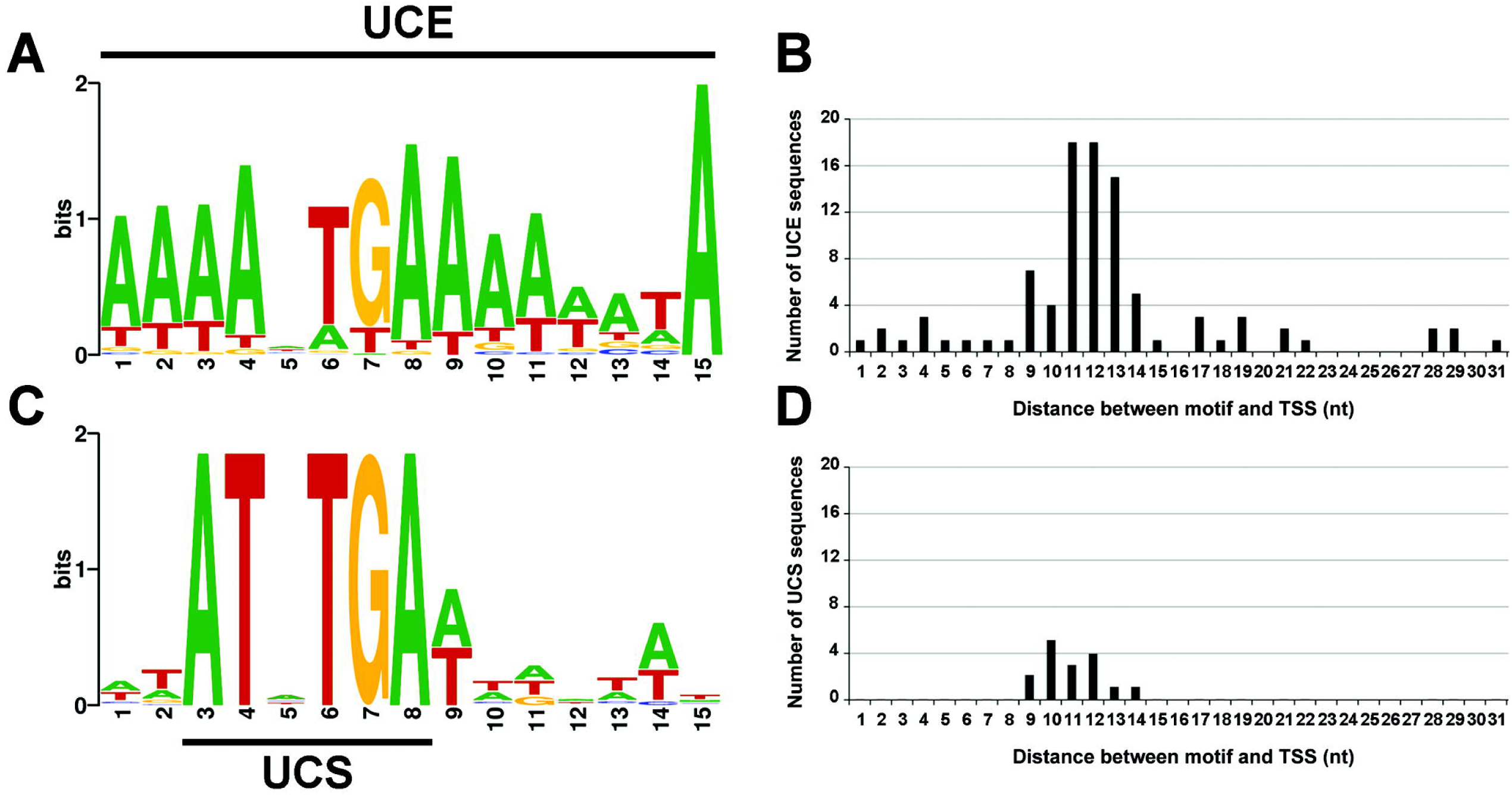
Upstream promoter elements of yeast linear plasmids show high sequence similarity to promoters of poxviral early genes. (**A**) Vaccinia virus early promoter consensus motif termed upstream control element (UCE) calculated from 84 sequences identified from genome-wide RNA-sequencing experiments in ref. (Yang et al. 2011). (**B**) Graph showing the number of UCE sequences as a function of their distances to the transcription start sites (TSS) of 84 ORFs as annotated in ref. (Yang et al. 2011). (**C**) Extended promoter consensus motif of pGKL plasmids preceding 15 ORFs. This motif contains the upstream conserved sequence (UCS) which is universal among yeast linear plasmids. (**D**) Graph showing the number of extended UCS sequences as a function of their distances to the TSS of 15 pGKL-encoded ORFs as determined in 5 RACE-PCR experiments. For promoters with a putative initiator region the first adenosine residue in the region was considered to be the TSS. For more information concerning promoter characterization see Supplementary Materials and Methods.

To conclude, it appears that promoters of poxviral early genes and linear plasmid genes are similar both with respect to their sequence and their spacing to the TSSs, implying a common origin.

## DISCUSSION

In this study we characterized the considerably underexplored transcription machinery of the yeast cytoplasmic linear double-stranded DNA plasmids. We used both experimental and bioinformatic approaches, and determined the composition and interactions of the transcription complex and presented a 3D model of its two main subunits. Further, we defined DNA sequences required for initiation and termination. For a model of the key aspects of transcription of the linear plasmid see Figure 9. Finally, our analyses provided evidence strongly suggesting that poxviruses and the yeast cytoplasmic DNA plasmids have a common origin.

**Figure 9.**
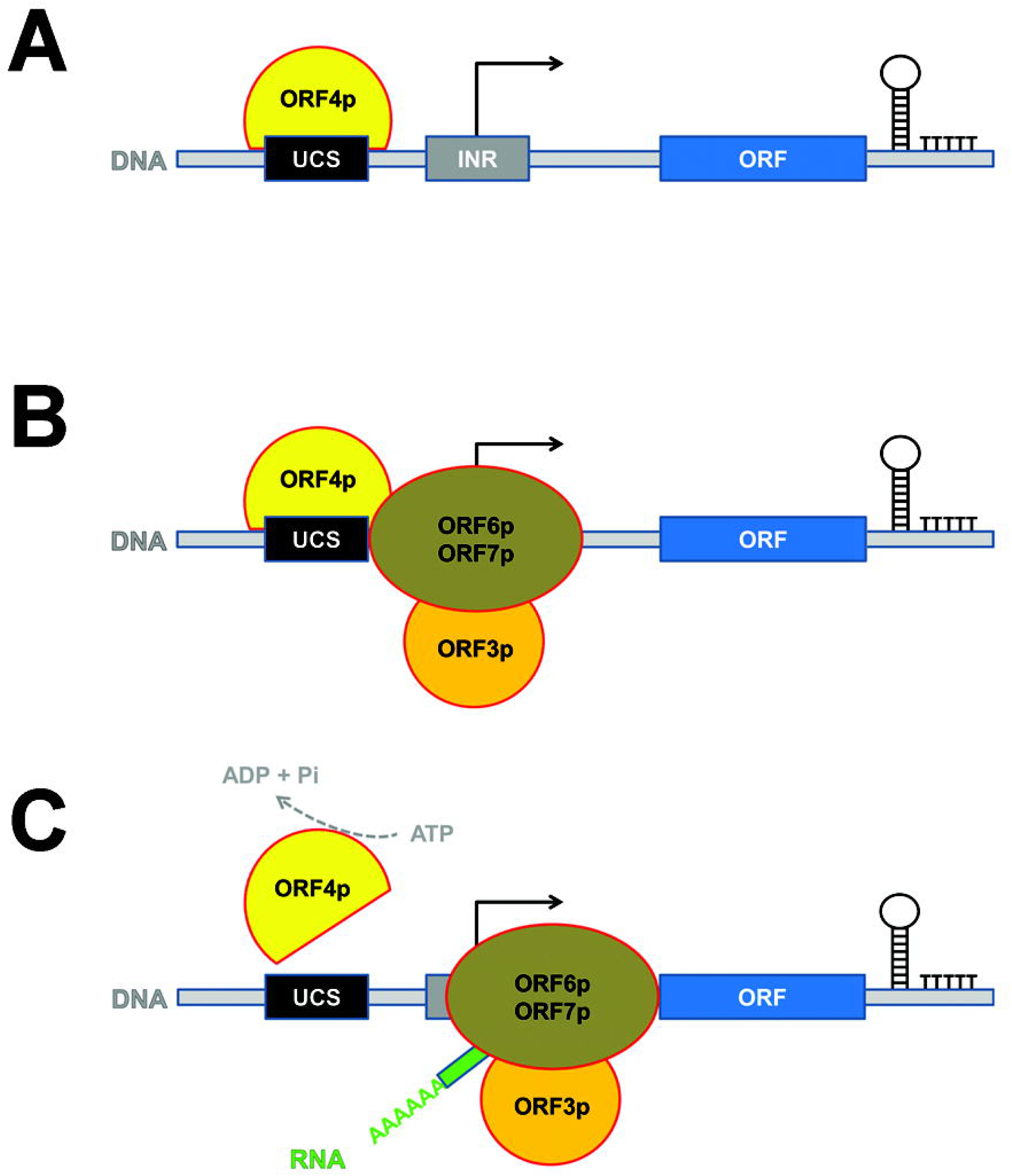
Model of the yeast linear plasmid transcription initiation and termination. (**A**) Putative helicase K2ORF4p (ORF4p, yellow) binds to the plasmid DNA, presumably to the upstream conserved sequence (UCS, black) which is related to the early promoter element of poxviruses. (**B**) K2ORF4p recruits the RNAP complex (ORF6p/ORF7p, brown) to the transcription initiation site, which usually contains the initiator region (INR, grey) responsible for RNAP slippage and subsequent 5’ mRNA polyadenylation. (**C**) ATP hydrolysis by K2ORF4p releases it from the transcription preinitiation complex to allow RNAP to escape from the initiation site and produce mRNA (RNA, green) containing a 5 end poly(A) leader. This RNA can be subsequently 5’ capped by the K2ORF3p viral-like mRNA capping enzyme (ORF3p, orange). Transcription termination most likely proceeds in a factor-independent manner that involves intrinsic terminators consisting of RNA stem loop structure(s) and 3’ terminal U-tract.

### Composition of the linear plasmid transcription machinery

Biochemical characterization of proteins encoded by yeast linear plasmids was shown to be challenging in the past. Expression of genes located on the pGKL plasmids seemed to be rather weak (Schründer and Meinhardt 1995; Schickel et al. 1996; Schründer et al. 1996). Also, it has been shown that expression of the K2ORF3p mRNA capping enzyme in routinely used *E. coli* systems was not possible, most likely due to the different codon usage dictated by the high AT content of plasmid genes (Tiggemann et al. 2001). Recently, it was shown that yeast nuclear expression of plasmid genes was impaired because the high AT content of plasmid genes led to RNA fragmentation (Kast et al. 2015). Accordingly, we failed to express recombinant K2ORF6p and K2ORF7p RNAP subunits in *E. coli, S. cerevisiae* and *K. lactis* expression systems (data not shown).

To overcome this difficulty, we prepared modified and double-modified pGKL plasmids expressing the transcription components with tags. Using co-immunoprecipitations followed by mass spectrometry and Western blotting we demonstrated that *in vivo* the transcription machinery core complex consisted of the two RNAP subunits (K2ORF6p, K2ORF7p) and the mRNA capping enzyme (K2ORF3p). This interaction was independent of the presence of nucleic acids. Subsequently, we showed that the putative helicase (K2ORF4p) associated less tightly with both the plasmid large RNAP subunit and the mRNA capping enzyme. This suggested the following molecular model of the plasmid transcription components interactions. First, we propose that K2ORF6p and K2ORF7p interact directly because aa residues of β’a15 and β’a16, which are known to participate in intramolecular *(Bacteria, Eukarya, Archaea)* or intermolecular *(Archaea)* bonds in multisubunit RNAPs, are present also in K2ORF6p and K2ORF7p as revealed by our *in silico* analysis. Second, we suggest that the mRNA capping enzyme interacts directly with the RNAP complex although it is yet to be determined whether it is with K2ORF6p and/or K2ORF7p. Cellular mRNA capping enzymes are known to interact with the C-terminal domain of RNAP II β’ homolog subunit (McCracken et al. 1997). However, a homologous C-terminal domain is present neither in Vaccinia virus nor in linear plasmid RNAPs. Nevertheless, in Vaccinia virus, the heterodimeric VTF mRNA capping enzyme interacts directly with the RNAP complex, and it is thought to be present both during transcription initiation and elongation (Luo et al. 1991; Hagler and Shuman 1992). By analogy, K2ORF3p may utilize a similar mode of interaction with its RNAP. Third, we propose that K2ORF4p binds to the RNAP core complex less tightly than the core subunits do between themselves. Vaccinia virus VETF and NPH-I helicases are known to interact with RNAP through an RNAP-associated protein of 94 kDa (RAP94), and this is specific for RNAP packaged in the virion (Ahn et al. 1994; Mohamed and Niles 2000; Yang and Moss 2009). VETF can interact with RNAP only in the presence of RAP94 (Yang and Moss 2009). However, there seem to be no RAP94 homologs outside poxviruses, which implies yet another undescribed mechanism of D6/D11 homologs binding to RNAP in other nucleo-cytoplasmic viruses. This might apply also to K2ORF4p of the yeast linear plasmids. Future studies will have to address the exact mode of K2ORF4p binding to the transcription complex. Finally, we found K2ORF4p to associate with pGKL-specific DNA *in vivo*. However, due to compact genomic organisation of pGKL plasmids precise mapping of the *in vivo* DNA binding sites of K2ORF4p using ChIP-seq would most likely turn out unsuccessful because resolution of the method would not match close spacing of the UCS promoter elements, which we presume might be the K2ORF4p binding sites. Our analysis indeed suggested close spacing of K2ORF4p binding sites *in vivo* because any pGKL region chosen for PCR amplification showed up to be enriched in anti-HA-K2ORF4p ChIP sample. Importantly, it seems that the transcription machinery of the yeast linear plasmids is remarkably if not entirely self-sufficient, because we did not find any cellular proteins to be specifically associated with the large plasmid RNAP subunit using mass spectrometry analysis.

### Transcription initiation

Our previous 5’ RACE-PCR experiments revealed short poly(A) leaders at the 5’ mRNA ends of most pGKL-encoded genes (manuscript submitted). These 5’ poly(A) leaders were heterogeneous in length among individual transcripts (1–21 adenosines per molecule) and not complementary to the template DNA. Non-template 5 poly(A) leaders are a characteristic feature of Vaccinia virus intermediate and late gene mRNAs that occur due to slippage of RNAP at the INR of the promoter (Bertholet et al. 1987; Schwer et al. 1987; Schwer and Stunnenberg 1988; Davison and Moss 1989). Some promoters of Vaccinia virus early genes containing the INR element can also produce mRNAs with short 5’ poly(A) leaders (Ahn et al. 1990; Yang et al. 2011). It has been shown that 5’ untranslated regions composed of 5 poly(A) leader sequences prior to start codon have a regulatory role in translation initiation (Shirokikh and Spirin 2008; Xia et al. 2011). Using bioinformatics, we detected a putative INR element in promoters of plasmid genes whose transcripts were 5’ polyadenylated. Using 5’ RACE-PCR and mutagenesis of the putative INR, we confirmed that the 5’ poly(A) leader of plasmid transcripts was associated with the identified INR element *in vivo*, and it was generated by the same mechanism as shown for Vaccinia virus postreplicative transcripts (Davison and Moss 1989). In Vaccinia virus, however, mutations in the INR of postreplicative promoters completely abrogate marker gene expression (Davison and Moss 1989; Baldick et al. 1992). This was clearly not the case for INR of pGKL promoters because expression of *G418^R^* was used for selection of clones containing recombinant plasmids. To assess possible role of INR alterations on plasmid gene expression, we prepared a strain (IFO1267_pRKL1–9) where we introduced TACCC mutations to the TAAAC INR of toxin subunit gene *K1ORF4*, and this strain showed reduced killer toxin production (Supplementary Figure S7).

### Transcription termination

We mapped the 3’ mRNA ends of all linear plasmid genes using 3’ RACE-PCR experiments. We identified 1–4 putative RNA stem loops close to the 3’ mRNA terminus of each ORF. Although the putative stem loops displayed relatively high values of Gibbs free energy (median of −7.5 kcal/mol), these values were comparable to the genome-wide predicted intrinsic terminators in *Mycoplasma hyopneumoniae* (median of −8.0 kcal/mol), an organism with a similarly high AT content (Fritsch et al. 2015). Further, RNA stem loops of bacterial intrinsic transcription terminators are usually followed by the typical 7–8 nt U-tract that promotes RNAP pausing at weak dA-rU DNA-RNA hybrid (Martin and Tinoco 1980; d’Aubenton Carafa et al. 1990; Gusarov and Nudler 1999). Interestingly, we detected T nucleotide enrichment in terminal 8 nt of plasmid-specific 3’ cDNA ends that corresponds to putative U-tract (Supplementary Figure S8). Importantly, we revealed a direct link between the putative RNA stem loop structure and the transcription termination pattern *in vivo*, suggesting an intrinsic transcription termination model for the yeast linear plasmids, similar to that in bacteria (reviewed in (Ray-Soni et al. 2016)). Even though we did not analyse the termination efficiency, we assume that transcription reads through at least some of the putative terminator sequences. Otherwise, functional expression of approximately half of the ORFs would not be possible due to the compact genomic organization inherent to pGKL plasmids. Future experiments will be required to understand this mechanism in more detail. Even though Vaccinia virus RNAP, presumably related to plasmid-specific RNAP, terminates transcription of early genes in a factor-dependent manner it was recently shown that RNA stem loops can influence both efficiency and location of transcription termination *in vitro* (Tate and Gollnick 2015). Therefore, proposed substantial reduction of plasmid-specific RNAP ancestor might have contributed to adaptation of this enzyme to transcription termination induced by RNA stem loops and possible loss of auxiliary factors required for this process.

### *In silico* 3D model of linear plasmid RNAP

Bioinformatic analysis of plasmid-specific RNAP proved to be challenging due to its unique reduced architecture and great evolutionary distance from other multisubunit RNAPs. From the 3D model it is evident that plasmid RNAP significantly differs from canonical RNAPs in several aspects:

(i) Almost the entire clamp structure element is absent. Only a basal portion of the clamp formed by βa15, βa16 and β’a20 conserved regions is maintained. The clamp is a mobile RNAP element and its closure is important for high stability and processivity of the enzyme. The clamp conformation is regulated by interaction of universally conserved elongation factors NusG and Spt4/5 with the clamp coiled-coil motif (Hirtreiter et al. 2010) – an element likely missing in plasmid RNAP. Therefore, it is highly unlikely that plasmid transcription machinery could use cellular Spt4/5 to increase processivity.

(ii) The lid and rudder elements are likely missing. The lid acts as a wedge to facilitate dislocation of RNA from the DNA-RNA hybrid molecule, and thereby maintains a constant size of the DNA-RNA hybrid between 7 to 10 base pairs (Vassylyev et al. 2007). However, it was shown, that lid-less bacterial and archaeal RNAPs were negatively affected when transcribing from ssDNA but not so much from dsDNA templates (Toulokhonov and Landick 2006; Naji et al. 2008). By analogy, RNA displacement from DNA-RNA hybrid molecule should not be affected by absence of the lid in plasmid RNAP transcribing double-stranded templates. The lid was also suggested to participate in bacterial intrinsic termination using stem loops (Vassylyev et al. 2007). However, bacterial RNAP without the lid was capable of intrinsic termination *in vitro* (Toulokhonov and Landick 2006). Therefore, we hypothesise that this structural feature is not crucial for intrinsic termination by plasmid RNAP *in vivo*. The rudder element interacts with the upstream edge of the DNA-RNA hybrid (Vassylyev et al. 2007). Experiments using bacterial RNAP with a deleted rudder reported defects in transcription initiation and less stable elongation complexes (Kuznedelov et al. 2002). This may correlate with the linear plasmid-specific termination of transcription.

(iii) The secondary-channel rim helices are missing. These helices are the binding sites for some transcription factors of multisubunit RNAPs, such as the transcription elongation factor TFIIS (Kettenberger et al. 2004). Therefore, it is highly unlikely that plasmid transcription machinery could use cellular TFIIS to overcome pause sites and increase proofreading.

### The evolutionary origin of the linear plasmids transcription machinery

A viral origin of the linear plasmid genes encoding the mRNA capping enzyme and the putative helicase was suggested previously (Jeske et al. 2007). However, the same origin for RNAP genes was not expected because previous hypotheses proposed that those genes originated from ancestral yeast RNAP genes (Jeske et al. 2007; Ruprich-Robert and Thuriaux 2010) or that they were ancient representatives of multisubunit RNAP diversification due to their simplified architecture (Iyer and Aravind 2012). However, no phylogenetic analysis was conducted to support the aforementioned hypotheses. Our results indicate that all the plasmid-specific transcription machinery components of the yeast linear plasmids have the same origin close to poxviruses. Poxviral RNAP, such as that of Vaccinia virus, lacks obvious α subunit homologs (Knutson and Broyles 2008), similarly to linear plasmid RNAP, and is more simplified than eukaryotic RNAPs. Therefore, a reduction of poxviral RNAP instead of yeast RNAP to give rise to the plasmid RNAP seems more plausible.

Our promoter analysis suggests that not just the plasmid RNAP, but also the plasmid promoters are related to nucleo-cytoplasmic viruses. We noticed sequence similarity between the UCE motif of Vaccinia virus early genes and the extended UCS motif of pGKL plasmids, as well as their similar location prior to the TSSs. Invariant G residue and several A residues in the minor groove were proposed to be the UCE nucleotides contacted by Vaccinia virus VETF helicase (Broyles et al. 1991). Presence of the invariant G residue and AT residues at other positions of the extended UCS motif suggests that K2ORF4p might contact UCS of the pGKL plasmids.

In conclusion, the transcription apparatus of the yeast linear plasmids has most likely an origin close to poxviruses and uses transcription initiation mechanisms similar to those used by poxviral genes. Unlike poxviruses, however, linear plasmids are beneficial for the cell, and this exemplifies the ability of the cell to domesticate originally harmful elements.

## MATERIALS AND METHODS

### Strains, plasmids and growth conditions

All of the strains used in this study are listed in Table 2. *Escherichia coli* cells were grown at 37°C in 2xTY medium which was supplemented with kanamycin (50 μg/ml) or ampicilin (100 μg/ml) for selection of transformants. Transformations of *E. coli* cells were performed by electroporation using Gene Pulser Xcell^TM^ (BIO-RAD). *K. lactis* cells were grown at 28°C in YPD medium which was supplemented with G418 (250 μg/ml) and/or hygromycin B (200 μg/ml) for selection of transformants. Transformations of *K. lactis* cells were performed using the one-step LiCl method (Gietz and Woods 2002) and followed by five-hour incubation in non-selective conditions immediately after transformation. For detailed descriptions of plasmids used in this study see Supplementary Table S1. Constructed pGKL plasmids were verified by PCR and subsequent sequencing of amplified products.

**Table 2.**
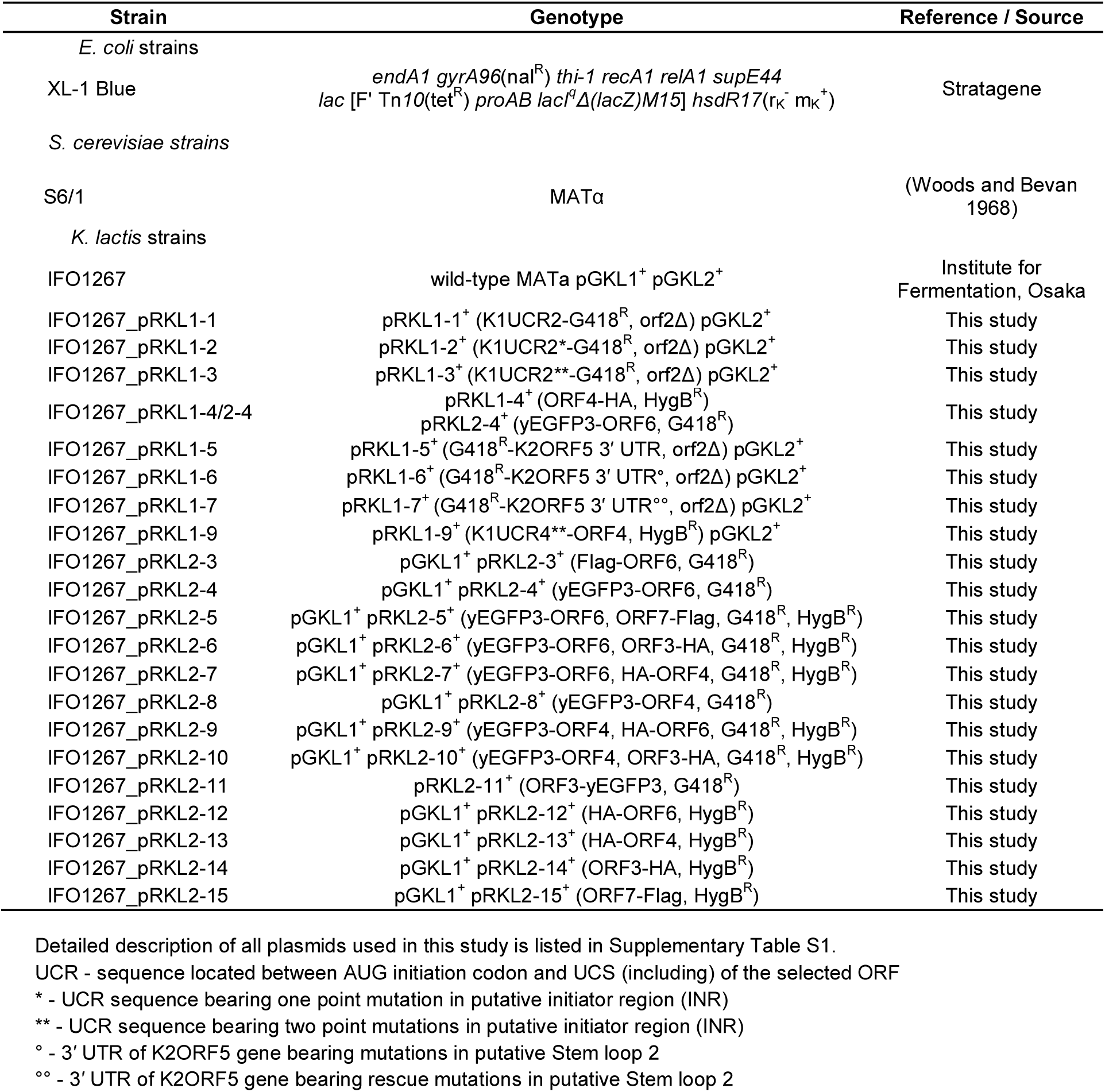
List of strains used in this study.

The nucleotide sequences of the primers used for construction, verification and sequencing of recombinant pGKL plasmids, and RACE-PCR amplification are listed in Supplementary Table S2. All polymerase chain reactions (PCRs) were performed using Taq DNA polymerase (Roche). PCRs for construction of recombinant pGKL plasmids were performed using mixture of Taq DNA polymerase (Roche) and Pwo DNA polymerase (Roche) in a 99:1 volume ratio, respectively.

### Modification of pGKL plasmids using homology recombination *in vivo*

*K. lactis* IFO1267 strain was transformed with PCR-generated fragment consisting of 5’ and 3’ ends homologous to the part of the pGKL plasmid to be modified and non-homologous part that introduced purification and/or detection tag (yEGFP3, HA-tag, Flag-tag) into plasmid-specific ORF together with a gene encoding resistance marker (G418 or hygromycin B) whose expression is driven by pGKL1-derived upstream control region (UCR, the sequence extending from the AUG initiation codon up to and including the UCS of the selected ORF). This type of construct was prepared by PCR or fusion PCR methodology (for details see Supplementary Table S3).

After PCR amplification and gel electrophoresis, corresponding fragments were purified using FavorPrep™ GEL/PCR Purification Kit (Favorgen) and used for transformation. After transformation, yeast cells were plated onto selective media and analyzed using agarose electrophoresis for the presence of the modified pGKL plasmid (for details see Supplementary Materials and Methods). Usually, it was possible to detect both, modified and wild-type target plasmid directly after transformation. Colonies containing both (modified and wild-type variants of the target pGKL plasmid) were selected and cultivated under selective conditions for approximately 60 generations and analyzed again using agarose electrophoresis. For subsequent analysis and preparation of double-modified pGKL plasmids, colonies containing only modified variant of the target pGKL plasmid were used. Absence of unmodified variant of the respective pGKL plasmid was also verified by PCR and subsequent electrophoresis.

### Co-immunoprecipitation and mass spectrometry

For the co-immunoprecipitation experiments, 100 ml of the yeast cells from the late exponential growth phase (OD_600_ = 4–8) were used. The cells were harvested with centrifugation, washed with distilled water and lysed in 2–3 ml of ice-cold GFP-wash buffer (10 mM Tris-HCl, pH 7.5, 150 mM NaCl, 0.5 mM EDTA) supplemented with 1 mM PMSF, cOmplete™ Mini protease inhibitors (Roche) and 0.45 mm glass beads using Mixer Mill MM 301 (Retsch) at a frequency of 30/s for 5 min. The glass beads and cell debris were pelleted with centrifugation at 8,000 g for 5 min at 4°C. The lysates were clarified with centrifugation at 20,000 g for 20 min at 4°C. Co-immunoprecipitations were performed using 25 μl of GFP-Trap^®^_A (Chromotek) beads with gentle mixing overnight at 4°C. Mock immunoprecipitations were performed using 25 μl of empty agarose beads (Chromotek) with gentle mixing overnight at 4°C. Bound proteins were washed three times in 1 ml of ice-cold GFP-wash buffer with gentle mixing for 5 min at 4°C. The immunoprecipitated complexes were dissolved in 60 μl of 2X sample loading buffer (0.1 M Tris-HCl, pH 6.8, 20% glycerol, 2% β-mercaptoethanol, 4% SDS and 0.04% bromophenol blue), incubated for 5 min at 95°C and subjected to SDS-PAGE. The gel was stained with Coomassie Brilliant Blue G-250 or silver. Gel lanes or bands of interest were excised from the gel, digested with trypsin (Shevchenko et al. 2006), and analysed by mass spectrometry. Identity of all detected proteins was also confirmed by MS/MS analysis. To test, whether certain co-immunoprecipitation is dependent on nucleic acids, the bound washed proteins were treated with 25U of Benzonase^®^ Nuclease (Novagen) in GFP-wash buffer for 30 min at 33°C and washed three times again, prior to SDS-PAGE analysis. Fraction of the bound washed proteins was taken and nucleic acids were eluted in 30 μl of TE buffer (10 mM Tris-HCl, pH 8.0, 1 mM EDTA), incubated for 5 min at 95°C, and 1 μl of the nucleic acids was used for detection of DNA using PCR amplification for 22–25 cycles and subsequent electrophoresis.

### Chromatin immunoprecipitation

For the chromatin immunoprecipitation experiments, 50 ml of the yeast cells from the late exponential growth phase (OD_600_ = 4–8) were used. The cells were cross-linked with formaldehyde (1% final concentration) added to the growing culture for 40 min at 28°C. The cells were harvested with centrifugation, washed with distilled water and lysed in 2 ml of ice-cold non-denaturing lysis buffer (50 mM Tris-HCl, pH 7.5, 300 mM NaCl, 5 mM EDTA, 1% Triton X-100, 0.02% sodium azide) supplemented with 1 mM PMSF, cOmplete™ Mini protease inhibitors (Roche) and 0.45 mm glass beads using Mixer Mill MM 301 (Retsch) at a frequency of 30/s for 5 min. The lysate was sonicated (Qsonica Ultrasonic Processor Q700, 50% amplitude) sixty times with 10 sec pulses. The glass beads and cell debris were pelleted with centrifugation at 8,000 g for 5 min at 4°C. The lysates were clarified with centrifugation at 20,000 g for 20 min at 4°C. The DNA fragments were immunoprecipitated using 30 μl of mouse monoclonal anti-HA HA-7 agarose (Sigma Aldrich) beads with gentle mixing overnight at 4°C. The beads were then washed once in 1 ml of ice-cold Wash buffer (20 mM Tris-HCl, pH 8.0, 150 mM NaCl, 2 mM EDTA, 1% Triton X-100, 0.1% SDS) supplemented with single stranded salmon sperm DNA (100 μg/ml; Roche), twice in 1 ml of ice-cold Wash buffer, and then once 1 ml of ice-cold Final wash buffer (Wash buffer containing 500 mM NaCl), each time with gentle mixing for 5 min at 4°C. Immunocomplexes were then eluted from the beads in 120 μl of Elution buffer (1% SDS, 100 mM NaHCO_3_) for 30 min at 37°C. Eluted immunocomplexes and 50 μl of the input clarified lysates were mixed with 400 μl of TBS (50 mM Tris-HCl, pH 7.5, 150 mM NaCl) supplemented with 5 μl of proteinase K (20 mg/ml; Sigma Aldrich), and the cross-linking was reversed by incubation for 5 hr at 65°C. The immunoprecipitated and input DNA was isolated by phenol-chloroform extraction followed by ethanol precipitation supplemented with 1 μl of linear polyacrylamide (25 mg/ml; Sigma Aldrich), and then used for PCR amplification for 25 cycles followed by electrophoresis. PCR amplifications were carried out on 1/30 of the chromatin immunoprecipitation (ChIP) and 1/1200 of the chromatin before immunoprecipitation (Input) using primers listed in Supplementary Table S2.

### Western blotting

SDS-PAGE gels were electroblotted onto Immun-Blot^®^ PVDF Membrane (BIO-RAD). The membranes were blocked in 5% non-fat dry milk (Hero) in a TBS-Tween buffer (50 mM Tris-HCl, pH 7.5, 150 mM NaCl and 0.5% Tween-20) and incubated with a primary antibody overnight at 4°C. After washing in TBS-Tween buffer and blocking with 5% non-fat milk, the membranes were incubated with a goat anti-mouse HRP-conjugated antibody (1:5,000 dilution; Santa Cruz Biotechnology). Finally, after washing in TBS-Tween buffer, the membranes were immersed in a luminol detection solution and the signal was detected using ImageQuant™ LAS 4000 (GE Healthcare). To confirm the expression of the target protein and the successful immunoprecipitation, a mouse monoclonal anti-Flag M2 antibody (1:1,000; Sigma Aldrich), mouse monoclonal anti-HA 6E2 antibody (1:1,000; Cell Signaling), and mouse monoclonal anti-GFP B-2 antibody (1:1,000; Santa Cruz Biotechnology) was used.

### RNA isolation, electrophoresis, reverse transcription, 5’ and 3’ RACE-PCR

25 ml of the yeast cells from the exponential growth phase (OD_600_ = 0.5–1) were quickly pelleted and frozen. Total yeast RNA was isolated by the hot acidic phenol procedure followed by ethanol precipitation (Lin et al. 1996). Remaining DNA was removed by DNA-*free*^TM^ Kit (Ambion). The quality of RNA was assessed by electrophoresis and UV spectrophotometry (Mašek et al. 2005).

In the case of 5’ RACE, subsequent reverse transcription was carried out as follows: 1 μg of total yeast RNA and 0.15 μg of random hexamer primers (Invitrogen) were used for cDNA synthesis using 100U of SuperScript^®^ III Reverse Transcriptase (Invitrogen). The cDNA was purified using High Pure PCR Product Purification Kit (Roche) and used for cDNA tailing using 800U of rTdT (Fermentas) and 0.5 mM dGTP (Roche) in 50 μl reaction for 30 min at 37°C with subsequent heat inactivation of rTdT for 10 min at 70°C. For PCR amplification of cDNA ends, 2.5 μl of the reaction mixture was used with olig2(dC)anchor primer and appropriate gene-specific primer for 35 cycles.

In the case of 3’ RACE, 1 μg of total yeast RNA was polycytidinylated using Poly(A) Tailing Kit (Applied Biosystems) and 2 mM CTP (Thermo Scientific) for 90 min at 37°C. Following reverse transcription was performed using 100U of SuperScript^®^ III Reverse Transcriptase (Invitrogen) and 1 μg of oligo(dG)anch2 primer. The cDNA was purified using High Pure PCR Product Purification Kit (Roche) and 2.5 μl of the purified cDNA was used for PCR amplification of cDNA ends with anch2 primer and appropriate gene-specific primer for 35 cycles.

In both RACE experiments, after PCR amplification and electrophoresis, obtained fragments were verified using restriction digestion and fragments exhibiting correct digestion pattern were gel-purified using FavorPrep™ GEL/PCR Purification Kit (Favorgen) and cloned to the pCR^TM^4-TOPO^®^ vector (Invitrogen). Vectors were transformed into *E. coli* XL-1 Blue cells (Stratagene), isolated using GenBond Plasmid FlexSpin Kit (Renogen Biolab) and sequenced using universal T7 promoter primer or T3 primer.

## ACKNOWLEDGEMENTS

We thank Natálie Suchánková and Jitka Vojáčková (Charles University, Czech Republic) for technical assistance. This work was supported by Czech Science Foundation (P305/12/G034), Charles University institutional project (SVV-260426), and ELIXIR CZ research infrastructure project (MEYS Grant No: LM2015047).

## COMPETING INTERESTS

The authors declare that no competing interests exist.

